# Molecular basis of the logical evolution of the novel coronavirus SARS-CoV-2: A comparative analysis

**DOI:** 10.1101/2020.12.03.409458

**Authors:** Abhisek Dwivedy, Krushna Chandra Murmu, Mohammed Ahmad, Punit Prasad, Bichitra Kumar Biswal, Palok Aich

## Abstract

A novel disease, COVID-19, is sweeping the world since end of 2019. While in many countries, the first wave is over, but the pandemic is going through its next phase with a significantly higher infectability. COVID-19 is caused by the novel Severe Acute Respiratory Syndrome Coronavirus 2 (SARS-CoV-2) that seems to be more infectious than any other previous human coronaviruses. To understand any unique traits of the virus that facilitate its entry into the host, we compared the published structures of the viral spike protein of SARS-CoV-2 with other known coronaviruses to determine the possible evolutionary pathway leading to the higher infectivity. The current report presents unique information regarding the amino acid residues that were a) conserved to maintain the binding with ACE2 (Angiotensin-converting enzyme 2), and b) substituted to confer an enhanced binding affinity and conformational flexibility to the SARS-CoV-2 spike protein. The present study provides novel insights into the evolutionary nature and molecular basis of higher infectability and perhaps the virulence of SARS-CoV-2.

## 1. Introduction

None of the recent outbreak like, SARS, HIV, Swine flu, could match the current pandemic COVID-19 except perhaps the flu pandemic that occurred over 100 years ago (Ashour et al., 2020). COVID-19 claimed around 54 million infections and over 1 million deaths globally while writing this report (WHO, 2020). SARS-CoV-2, unlike many other viruses, can be spread by asymptomatic individuals (Andersen et al., 2020; Ashour et al., 2020). Elucidating the molecular and cellular bases of the viral infection would enhance the understanding of the virulence of the virus.

Besides major damage to the respiratory system following SARS-CoV-2 infection, other important associations of the disease are neurological defects (overtly loss of taste and renal failure, coagulopathy and vascular disease along with many other conditions (Jin et al., 2020; Rothan and Byrareddy, 2020). A genome-wide association study linked an increased susceptibility to the COVID-19 in patients with blood group A and in males (Zhao et al., 2020). Among co-morbidities, hypertension and diabetes are the main concerns. In addition, the virus itself presents some intrinsic yet unknown features to enhance its virulence like-a) proof-reading mechanism(s) to protect itself from external adverse effects and agents; b) a larger genome, thrice that of HCV and double that of influenza virus (Benvenuto et al., These along with many other molecular features of SARS-CoV-2 have made the specificity and affinity of its spike protein, for ACE2 of the human host, significantly higher (Walls et 2020). The higher affinity, dynamic rearrangement, and specificity of the SARS-CoV-2 spike protein for ACE2 are among the key factors that might have made the virus more virulent (Yan et al., 2020). The pertinent question is how it acquired such potential and precise machinery within a short span, following the SARS-CoV and MERS-CoV that took place in 2003 and 2012. We, therefore, attempted to understand the evolution of coronavirus of various kinds, with special emphasis on SARS-CoV, MERS-CoV and SARS-CoV-2 to determine cues on the evolutionary dynamics that have enhanced the virulence and infectivity of SARS-CoV-2. We have analyzed the amino acid sequences of Spike proteins of 45 relevant coronaviruses and the structural features of select ones to understand the major differences that might explain the increased binding efficiency of the SARS-CoV-2 spike proteins to human ACE2. The Spike proteins from coronaviruses into two distinct fragments, S1 and S2. Fragment S1 is involved in recognition of host cell surface receptors, and the fragment S2 is involved in generation of the pre-fusion complex. Fragment S1 is comprised of two major domains-N-terminal (NTD) and C-terminal (CTD) domains (Li, 2016). Collectively, NTD and CTD are also known as the receptor binding domain (RBD). The CTD interacts with molecules like ACE2 and CD26, in case of SARS-CoV/CoV-2 and MERS/Bat-CoV, respectively. The NTD is known to recognize sugar containing molecules and cell adhesion molecules (Li, 2016; Sun et al., 2020). The physiological state of the Spike proteins is comprised of a homo-trimer with a central three-fold symmetry with the three S1 fragments sitting atop the respective membrane anchored S2 fragments (Figure 1A) (Li, 2016). We validated the protein structural data by analyzing the differences in the coding nucleotide sequences. The results showed the plausible mutations that act as the driving force in the natural selection of SARS-CoV-2.

**Figure 1:**
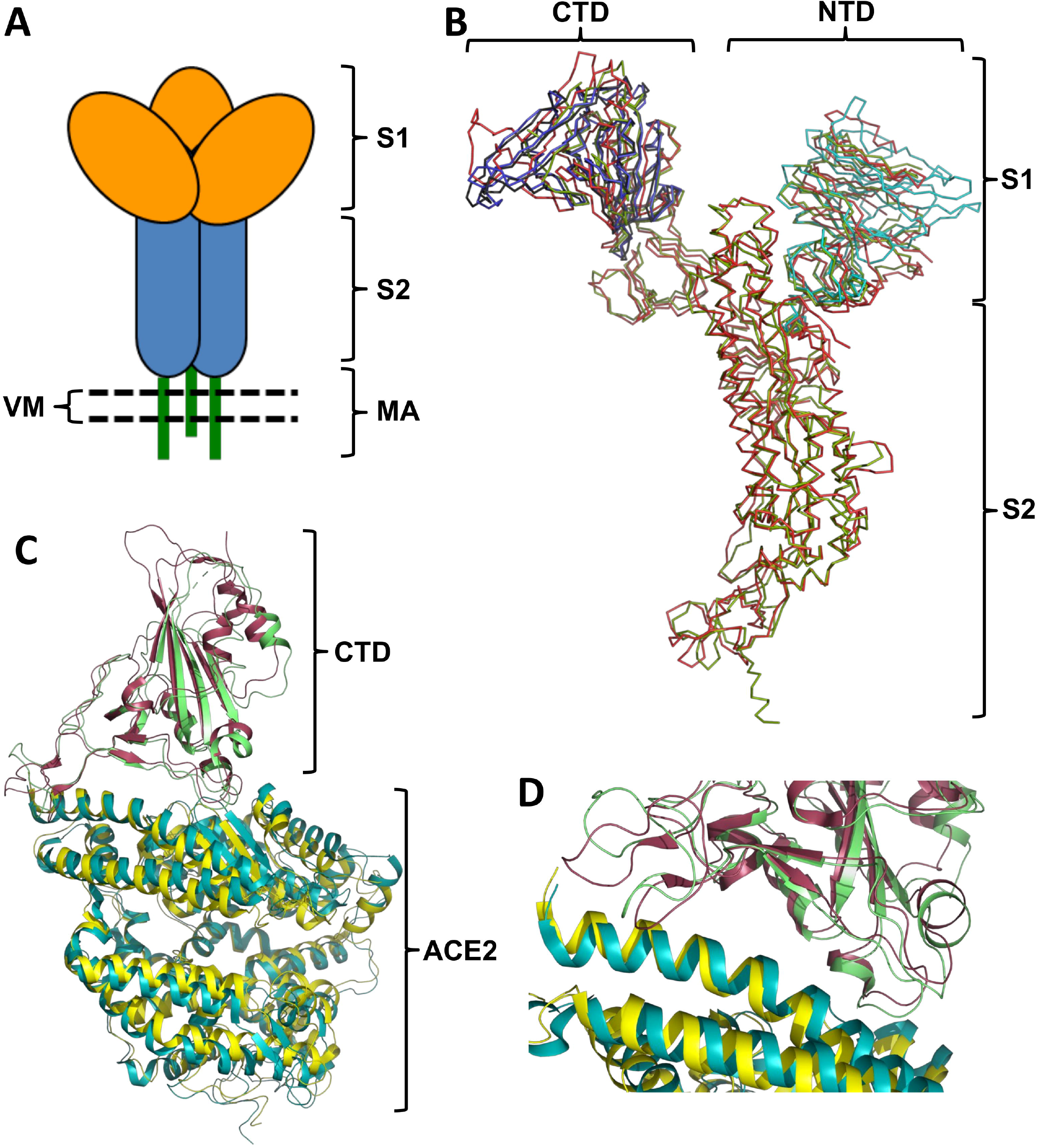
The 3-D structure comparison of Spike proteins of SARS-CoV and SARS-CoV2 and their interaction interfaces with human ACE2. **(A)** A cartoon representation of the Spike protein trimer assembly depicting the position of sub-units S1 and S2 and the membrane anchor (MA) region on the viral membrane (VM). **(B)** A superimposition of the Cα chains for the complete spike protein from SARS-CoV2 (lime-green); complete spike protein from SARS-CoV (tv-red); NTD of spike protein from Bov-CoV (cyan); CTD of spike protein from MERS-CoV (dark-grey) and CTD of spike protein from Bat-CoV (tv-blue). Of note, the CTD of the SARS-CoV2 is more compact as compared to SARS-CoV, MERS-CoV and BAT-CoV. **(C)** A superimposition of the NTD of the SARS-CoV2 and SARS-CoV bound to the human ACE2 – Cα chain in left panel and secondary structures in right panel. (SARS-CoV2 in raspberry, CoV2 bound ACE2 in cyan; SARS-CoV in tv-green, CoV bound ACE2 in yellow). **(D)** A superimposition of the interacting regions of the NTD of the SARS-CoV2 and SARS-CoV and the human ACE2. (SARS-CoV2 in raspberry, CoV2 bound ACE2 receptor in cyan; SARS-CoV in tv-green, CoV bound ACE2 in yellow).

## 2. Materials and Methods

### 2.1 Protein Sequence and Structure analysis

The sequences of 45 Coronavirus Spike proteins were retrieved from the SwissProt database. The details of the sequences are presented in Table S1. The sequences were aligned using Clustal Omega (Sievers and Higgins, 2014) and a maximum likelihood phylogenetic tree was generated using the NEXUS algorithm (Giribet, 2005). The sequence alignments were represented using Espript 3.0 (Gouet et al., 2005).

The 3-D structures of Spike Proteins (native proteins, Fragment S1-C-terminal domains in complex with host receptors) from SARS-CoV-2 (IDs-6VYB, 6M0J), SARS-CoV (IDs-5XLR, 3SCI), MERS-CoV (ID-4L72), Bat Coronavirus HKU14 (ID-4QZV) and Bovine Coronavirus (ID-4H14) were retrieved from the Protein Data Bank. Multiple structure alignments were performed at the POSA web server (Li et al., 2014) using both flexible and rigid body algorithms. Two structure alignments were performed at the FATCAT web server (Veeramalai et al., 2008) using both flexible and rigid body algorithms. Complex of Fragment S1-C-terminal domains with the ACE2 binding domain were generated using the ZDOCK web server (Pierce et al., 2014) and the complexes generated were refined using the GalaxyRefineComplex web server (Heo et al., 2016). Interaction and binding properties of Spike proteins’ C-terminal domains with host receptor proteins were predicted using the PRODIGY web server (Xue et al., 2016). Surface area of interactions between Spike proteins’ C-terminal domains with host receptor proteins were determined using the InterProSurf (Negi et al., 2007) and the PISA (Baskaran et al., 2014) web servers. *In silico* alanine scanning mutagenesis for the protein-protein complexes were performed at the DrugScore^PPI^ (Krüger and Gohlke, 2010), the SpotOn (Moreira et al., 2017) and the mCSM-PPI2 web servers (Rodrigues et al., 2019). Pymol 2.3 was used for structural visualizations (DeLano, 2020).

### 2.2 Nucleotide sequence analysis

45 nucleotide sequences of the coronavirus Spike proteins were retrieved from NCBI nucleotide database. The details of the sequences are presented in Table S1. The sequences were aligned using MEGA X software (Kumar et al., 2018) with MUSCLE (Edgar, 2004) as the alignment algorithm using the default parameters. Post alignment, a distance matrix was calculated from the aligned output followed by both neighbour joining (NJ) and unweighted pair group method with arithmetic mean (UPGMA) methods. Following this, distance matrix calculator for maximum Parsimony and Maximum likelihood analyses were performed. The phylogenetic trees were generated using the R-package “ggtree” of Bioconductor including the genera and host of the respective coronavirus (Yu et al., 2017). For maximum likelihood we estimated the best model using the modelTest function from the “phangorn package” (Posada and Crandall, 1998). GTR+G+I was selected as the model to perform Maximum likelihood phylogenetic tree with 100 iterations. The sequence alignments were represented using Espript 3.0. The ancestry and substitution analysis were performed using MEGA X.

### 2.3 Dendrogram comparison analysis

Aligned amino acid and nucleotide sequences were assigned the same names in both the alignments for comparison. The phylogenetic distances were calculated for UPGMA using the “phangorn library” (Schliep, 2011). The best model was calculated for both nucleotide and amino acid sequences using the “modelTest” function where the “Akaike Information Criterion” (Ingram and Mahler, 2013) was applied to determine the best model for both trees and then each tree was converted into a dendrogram using “as.dendrogram” function. These dendrograms were further taken for dendrogram comparison using the “dendextend tanglegram function” (Galili, 2015). Tree distance was calculated using “treedist” function (Smith, 2020).

### 2.4 Synonymous and non-synonymous mutation analysis

For synonymous and non-synonymous mutation analysis, the 45 nucleotide sequence files in which the headers were labelled same as the respective amino acid sequences were used. Using the reverse align function the nucleotide sequences were reverse aligned by seqinr library (Charif and Lobry, 2007). The webserver RevTrans 1.4 (Wernersson and Pedersen, 2003) was used to generate amino acid/codon based alignment of nucleotide sequences. The codon aligned sequences were used to determine the synonymous (K_a_) and non-synonymous (K_s_) substitutions. By using these values, we evaluated the *dN, dS* and *dN/dS* value matrices. These matrices were further used for visualization of the sequence clustering using heatmaps. “Ward D2” was used as the clustering algorithm to cluster the samples into the genera and primary hosts of the respective virus.

## 3. Results

### 3.1

SARS-CoV-2 has both structural and functional similarities with the previous human coronaviruses but with much higher infectivity and lower morbidity (Walls et al., 2020). It is further noted that besides the similar host recognition molecules, SARS-CoV-2 has a higher affinity and a tighter binding with the human cellular receptor (Wrapp et al., 2020; Yan et al., 2020). These two features of SARS-CoV-2 prompted us to ask an important and obvious question-how and when did the novel coronavirus emerge to be a distinct lineage in terms of its enhanced infectability? What are the molecular markers that can be analyzed to understand the molecular basis of the stronger affinity and higher infectability? In the current report, we investigated these questions by-a) comparing the protein structures of spike proteins from SARS-CoV, MERS-CoV, SARS-CoV-2 and other related Bat-CoVs; b) establishing the similarity and differences in major amino acid residues to understand the higher affinity of SARS-CoV-2 towards ACE2 binding compared to other coronaviruses; and c) comparing the nucleotide and amino acid sequences of spike proteins to estimate the most probable evolutionary trend.

#### 3.1.1. A comparison both at the sequence and structure levels of the coronavirus Spike proteins

A multiple sequence alignment of 45 experimentally verified Spike proteins sequences from several species of coronaviruses showed a significant difference in the conservation status of the two fragments S1 and S2 (Supplementary File S1). The fragment S2 (aa 662-1272 for SARS-CoV-2) exhibited a significantly higher sequence conservation, with 76 amino acids strictly conserved across species. However, the fragment S1 (aa1-aa661 for SARS-CoV-2) exhibited an unusually low conservation with only 10 strictly conserved residues, possibly attributed to the wide repertoire of the host receptor molecules recognized by this domain (Li, 2016). The amino acid cysteine displayed the highest conservation across all sequences, at 16 different positions across the length of the sequences. Spike proteins form various inter-and intra-molecular di-sulphide bonds in order to stabilize the core monomeric structure as well the physiological homo-trimeric form (Li, 2016). A recent study by Wang et al. on the crystal structure of the SARS-CoV-2 Spike Protein CTD in complex with human ACE2, established the residues of CTD (K417, G446, Y449, Y453, L455, F456, Y473, A475, G476, E484, F486, N487, Y489, F490, Q493, G496, Q498, T500, N501, G502, and Y505) that interact with the human ACE2 (Figure S1) (Wang et al., 2020). Of these 21 residues, only 8 residues (Y436, Y440, N474, Y475, Y484, T486, G488, and Y491) are conserved in the SARS-CoV CTD.

In order to investigate the evolutionary divergence of the Spike protein, we generated an evolutionary tree (Figure S2). Interestingly, of the seven known human coronaviruses, the three coronavirus species associated with higher infectivity and morbidities, i.e., SARS-CoV, MERS-CoV and SARS-CoV-2 formed a distinct evolutionary cluster. Notably, the other members of these clusters were overtly the bat coronavirus species. Such a specific clustering suggests a possible co-evolution in the Spike proteins of humans and bat coronaviruses that led to the association with more severe infections.

#### 3.1.2. Comparison of the 3-D structures of the Spike proteins reveals residues critical for ACE2 binding

This association of functional features prompted an investigation into the structural similarities of the spike proteins from aforementioned human coronaviruses associated with high fatality and bat coronavirus. While the spike proteins SARS-CoV-2 and SARS-CoV displayed an overall conservation of the protein structure (Figure S3 A), the CTD domain of the SARS-CoV-2 is relatively compact and contains, unlike SARS-CoV, an extended long loop. (Figure S3 B & C). Notably, most of the β-strands of the central sheet in SARS-CoV-2 CTD are longer in size than that of the corresponding sheet in SARS-CoV. The scenario is however much different in other coronaviruses. Particularly, the CTD of Bat-CoV and MERS-CoV are larger with an anti-parallel β-sheet replacing the loop like structures of the ACE2 recognizing region of the CTD of SARS-CoV-2 (Figure S4). This comparative structural analysis hinted at a divergent evolution of the CTDs into two independent lineages-a) MERS-CoV and b) SARS-CoV-2. We also found that the core structure of the NTDs of the SARS-CoV-2 spike protein and the lectin binding NTD of Bov-CoV (Bovine Coronavirus) spike protein are largely similar (Figure S5). Taken together, the results suggest that the evolution of the host receptor recognizing domain in the coronavirus spike proteins are more local in nature while the global architecture demonstrate significant conservation (Figure 1B).

To explore these subtle evolutionary changes, we probed into the local architecture of the interfaces of SARS-CoV-2 CTD/hACE2 and SARS-CoV CTD/hACE2 complexes reported in the cryo-EM determined structures (Figure 1C) (Wrapp et al., 2020). Our analyses led to important findings that could help not only understanding the evolution and origin of the SARS-CoV-2 but also will help in developing potential intervention. Apart from the two small β-strands present in the SARS-CoV-2 CTD, the interfaces in the complexes were primarily lined up with loop like structures from the CTD of the spike protein and the N-terminal helix of the ACE2 (Figure 1D). It is important to note that SARS-CoV-2 CTD has 21 residues that interact with ACE2 N-terminal helix, while the SARS-CoV CTD has only 17 interacting residues. A closer inspection of the amino acid residues, involved in the interactions, suggested that residues Y453, Y473, G476 and F486 from SARS-CoV-2 CTD were crucial towards providing a stronger interaction with ACE2, with no identical residues from SARS-CoV in the respective molecular environment (Figure 2A). In order to determine any evolutionary correlation, the MERS-CoV and Bat-CoV CTDs were docked onto N-terminal helix region of the ACE2 followed by *in silico* energy minimization of the complexes. The MERS-CoV and Bat-CoV CTDs exhibited 18 and 19 residues interacting with ACE2 N-terminal helix, respectively (Figure 2B & 2C). Notably, the Y453 of SARS-CoV-2 superimposed with the identical interacting residues Y499 and Y503 from MERS-CoV and Bat-CoV CTDs, respectively in the molecular microenvironment (Figure 2B & 2C). In order to understand the contribution of each interacting residue of the CTDs in ACE2 binding, *in silico* alanine scanning mutagenesis analysis was performed. While Y453 of SARS-CoV-2 contributed 2.018 kcal mol^-1^, F486 contributed 3.01 kcal mol^-1^ to the interaction. Interestingly, Y499 and Y503 of MERS-CoV and Bat-CoV CTDs contributed significantly higher to their respective interactions-2.877 and 3.017 kcal mol^-1^, respectively (Figure 2D). Also, a significant rise was observed in the dissociation constants (K_d_) of binding with the ACE2 following alanine mutations of the aforementioned residues (Figure 2D). These energy values suggest the spatial conservation of this tyrosine residue in the CTD of SARS-CoV-2 being key to a stronger ACE2 binding, which is completely absent in the CTD of SARS-CoV.

**Figure 2:**
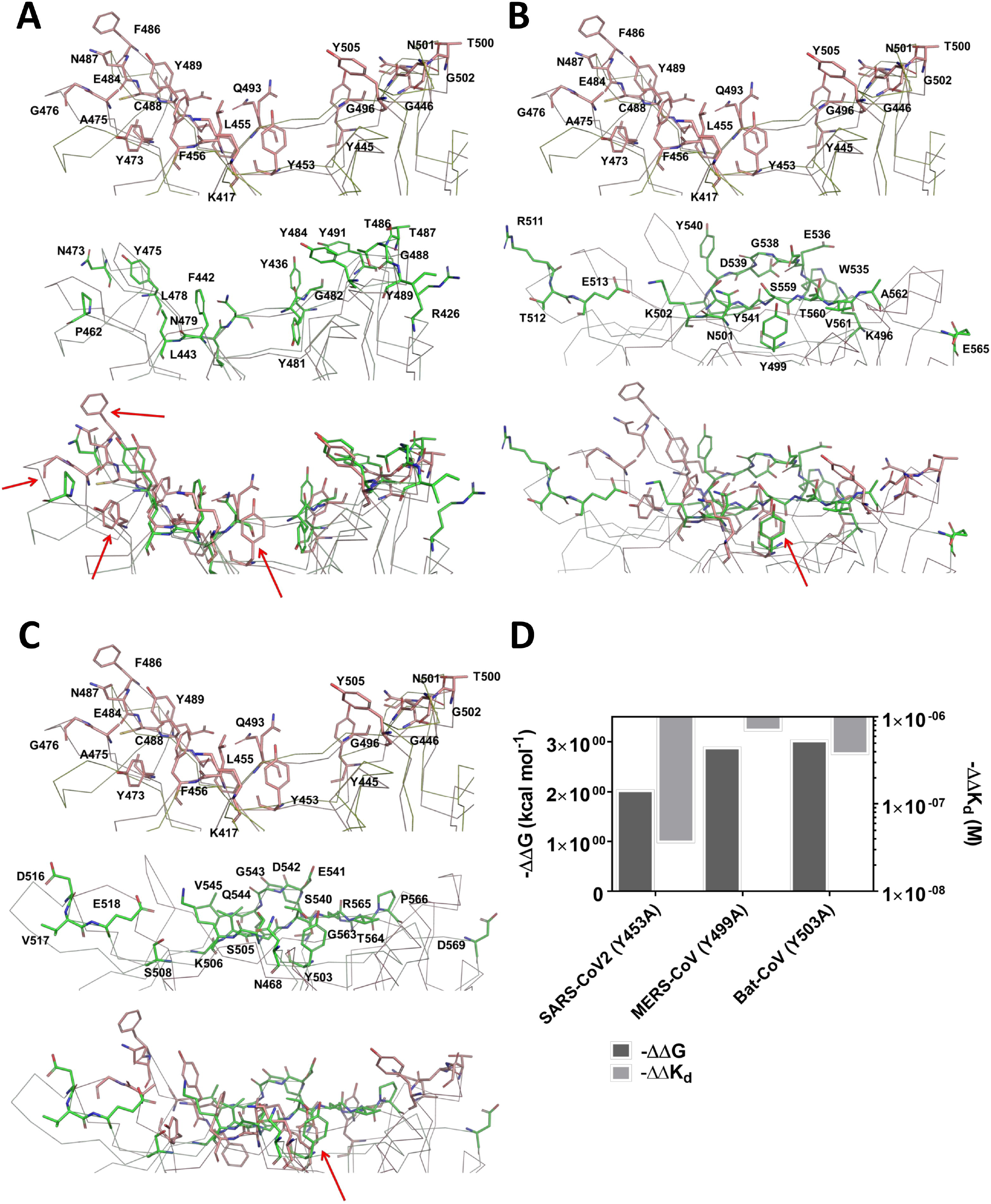
Comparison of the amino acid residues of the Spike proteins of SARS-CoV2, SARS-CoV, MERS-CoV and Bat-CoV involved in interactions with human ACE2. **(A)** Superimposition of the ACE2 binding residues of the CTD of SARS-CoV2 (raspberry) and SARS-CoV (pale green), with residues from each highlighted in sticks. (Crucial interacting residues from CoV2 marked in red arrows). **(B)** Superimposition of the ACE2 binding residues of the CTD of SARS-CoV2 (raspberry) and MERS-CoV (pale green), with residues from each highlighted in sticks. (Crucial interacting residue from CoV2 with identical superimposed residue from MERS-CoV marked in red arrow). **(C)** Superimposition of the ACE2 binding residues of the CTD of SARS-CoV2 (raspberry) and Bat-CoV (pale green), with residues from each highlighted in sticks. (Crucial interacting residue from CoV2 with identical superimposed residue from MERS-CoV marked in red arrow). **(D)** *In silico* analine scanning shows the significance of Y453 from SARS-CoV2 CTD and its spatial conservation in MERS-CoV and Bat-CoV.

A comparison of the *in silico* binding properties of the four aforementioned CTDs with the ACE2 revealed that despite a higher K_d_ for SARS-CoV-2, there was a significant decrease in the surface area of interaction, suggesting a higher specificity of interaction between residues of the CTD and the ACE2 N-terminal helix (Table 1). It is also worth noticing that while the interacting residues are widely spread across the interacting surface of the SARS-CoV CTD. However, for SARS-CoV-2 CTD the interactions localize on the far ends of the interacting surface (Figure S6 A-D). This phenomenon is crucial as the central region of the interacting surface is primarily comprised of uncharged residues that arch away from the N-terminal helix of the ACE2 in both SARS-CoV-2 and SARS-CoV CTDs. In a stark cont-rast, majority of the interacting residues in MERS-CoV and Bat-CoV CTDs were localized in the central region of the interacting surfaces. This observation further hinted at a divergent evolution, resulting in the formation of the β-sheet protruding out of the region in case of MERS-CoV and Bat-CoV CTDs, as opposed to loop like structures in SARS-CoV-2 and SARS-CoV CTDs.

#### 3.1.3. Maximum Likelihood and Parsimony analyses of the Spike protein nucleotide sequences at a glance

The nucleotide sequences of the spike proteins were aligned. The results were plotted for both maximum parsimony (Figure S7A) and maximum likelihood (Figure S7B) trees. The amino acid phylogenetic analysis showed a significant cluster overlapping among SARS-CoV (P59594), MERS-CoV (K9N5Q8) and SARS-CoV-2 (P0DTC2). These coronaviruses already established high morbidities in humans. These groups of coronaviruses cluster separately with several Bat-CoVs as observed in the amino acids sequences phylogenetic tree. Comparison of the Parsimony and Maximum likelihood analyses and the structural data together established the facts that SARS-CoV, Bat-CoV, MERS-CoV and SARS-CoV-2 belong to a similar functional, structural and evolutionary cluster. A similar pattern was observed in case of nucleotide phylogenetic analysis where the aforementioned coronaviruses form a distinct cluster. A co-phylogenetics analysis was performed to compare the nucleotide and amino acid sequence dendrograms (Figure 3). The trees exhibited an entanglement score of 0.22, with a topological distance of 1.548986. The treedist analysis suggested a symmetric difference score of 60.000000; branch score difference of 1.764349; path difference of 166.679333; and quadratic path difference of 20.924544, suggesting that tree are near identical despite the noticeable differences in the evolutionary lineages. The Baker’s Gamma correlation coefficient (Baker, 1974) was calculated to be 0.4489648 and the Cophenetic correlation (Lapointe and Legendre, 1995) between amino and nucleotide trees the value was found 0.8245775. These values suggest a significant similarity between the two trees. Taken together, these analyses suggest that majority of the changes observed in the nucleotide sequences were pronounced in the difference in amino acids, with a negligible codon bias.

**Figure 3:**
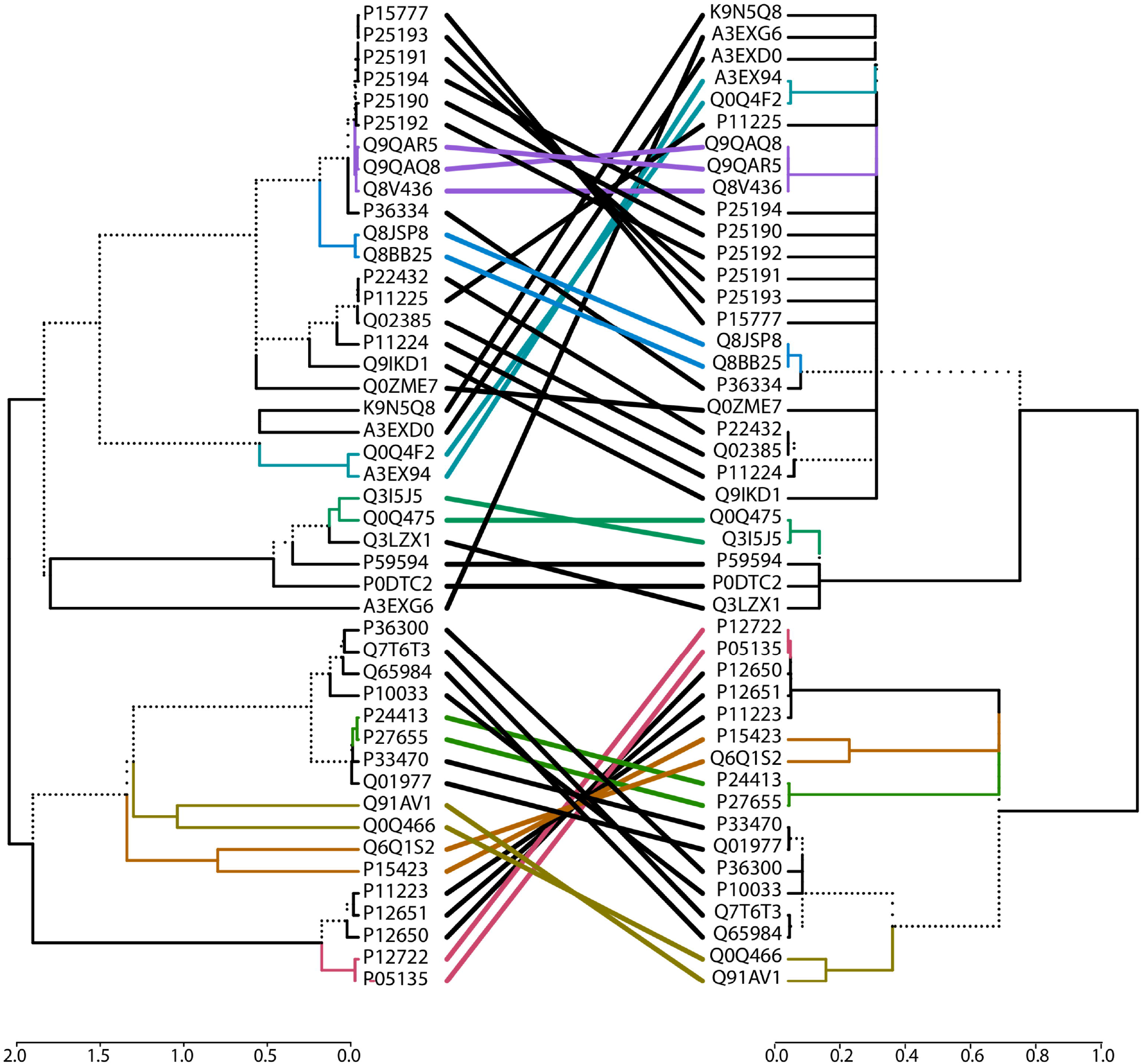
A tanglegram comparsion for the Spike protein amino acids and nucleotide phylogenies. With an entanglement score of 0.22, the phylogenies suggest a strong concordance, suggesting that codon usage has negligible impact on the divergence of Spike proteins. (Dashed branches differ between both phylogenies and coloured clades are identical to both trees. Amino acid phylogeny in left panel and row matched Nucleotide phylogeny in the right panel).

#### 3.1.4 Estimating the selection pressure on Spike proteins

We sought to investigate the possible selection pressure on the Spike proteins genes, particularly targeting the SARS-CoV-2 Spike protein. The numbers of non-synonymous substitutions *(dN)* between species, and the number of synonymous substitutions *(dS)* between species were calculated and used to determine the ratio of *dN* and *dS* (Rocha et al., 2006). A higher *dN* is associated with a positive selection, suggesting that associated mutations not only cause increased fitness but also indicate a recent divergence in species. Concurrently, a higher *dS* is indicative of a purifying selection, that remove deleterious mutations which reduced fitness. We estimated the *dN* and *dS* values for the set of 45 Spike proteins’ polypeptide sequences. While the *dN* values varied between −6 to 2, a significant proportion of the clusters depicted value greater than 1, suggesting higher number of non-synonymous substitutions across the spike protein sequences (Figure S8). More importantly, the *dS* values varied from −4 to 4 (Figure S9) with a majority of the values being less than 1. Such a distribution of *dS* values indicated that purifying selection in Spike proteins is limited to a small section of the coronaviruses. In stark contrast to the overall higher *dN* scores, it was strikingly low within SARS-CoV-2, SARS-CoV, and Bat-CoV suggesting a negligible positive selection. In concurrence, the *dS* scores within the aforementioned Spike proteins, suggested a possibly weak purifying selection. However, the comparison of MERS-CoV with SARS-CoV-2, SARS-CoV, and Bat-CoV suggested a strong positive selection as was evident from the high *dN* and the low *dS* values. In order to further understand the nature of selection pressures on Spike protein, the ratios of *dN* to *dS* were estimated. The ratios revealed a distribution range from −4 to 4 and presented an interesting scenario (Figure 4), suggesting a mild purifying selection driving the Spike protein evolution in SARS-CoV-2 and a strong positive selection in case of MERS-CoV. In addition, the comparison also suggested that the divergence of MERS-CoV could be an evolutionarily recent event while the evolution of SARS-CoV-2 might have occurred over a comparatively longer time span.

**Figure 4:**
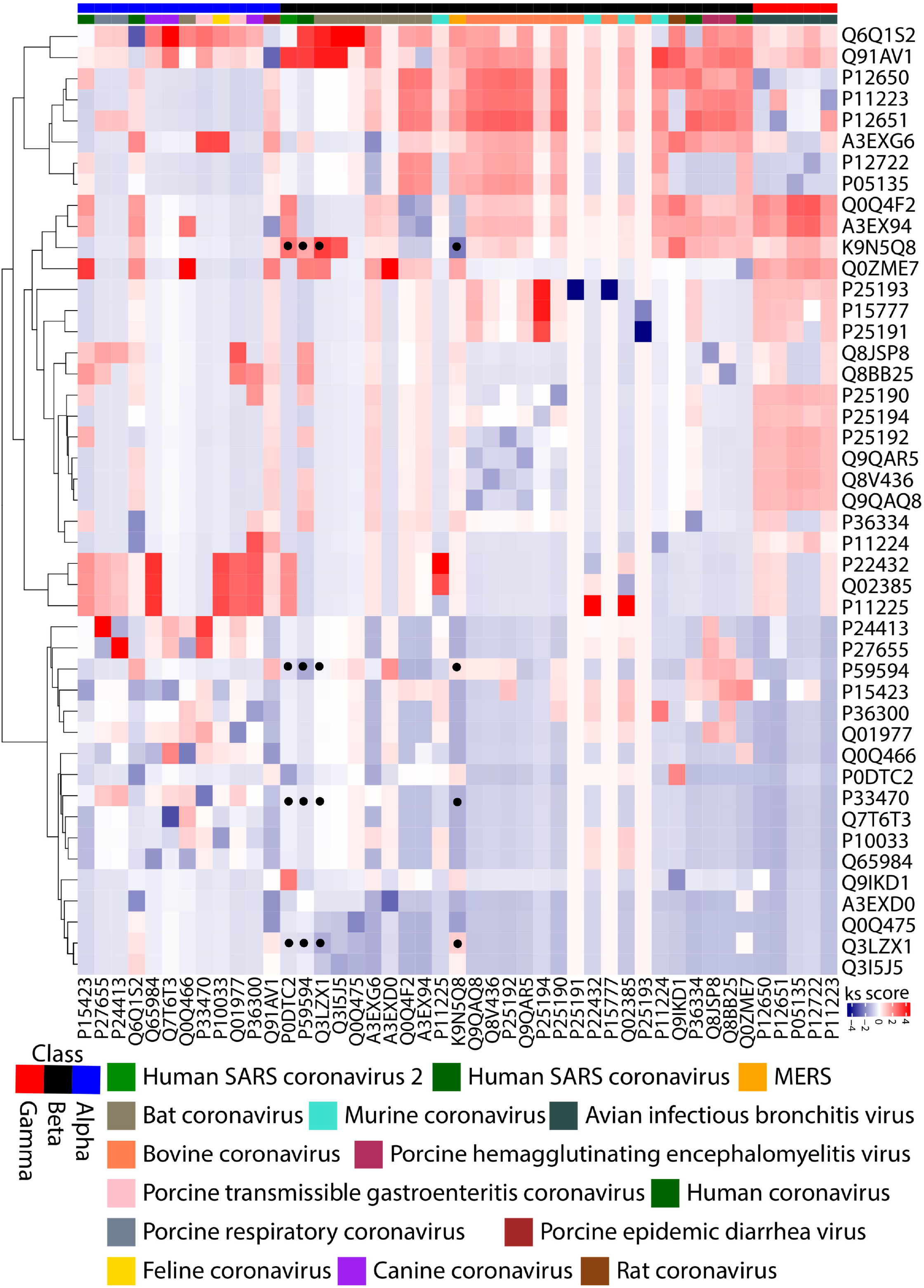
A heat map demonstration of the clustering of the *dN* – *dS* ratio for Spike protein coding ORFs. The values suggest a mild purifying selection for SARS-CoV2 and a strong positive selection for MERS-CoV. (Values within SARS-CoV2, SARS-CoV, MERS-CoV and Bat-CoV are marked with a black dot).

#### 3.1.5 A comparison of the amino acids and nucleotide sequences of the coronavirus Spike proteins

To better understand the aforementioned evolutionary conundrum, we closely examined the protein (Figure S10) and nucleotide (Figure S11) sequence of CTDs of five Spike protein sequences – SARS-CoV-2, SARS-CoV, MERS-CoV, and Bat-CoV (all belonging to the sub-genus Sarbecovirus) (Table S2). The CTDs of these Spike proteins exhibited 25 conserved residues. The CTDs of MERS-CoV and Bat-CoV are evidently longer and shorter, respectively in comparison to the SARS-CoV and SARS-CoV-2 CTDs. The theoretical pI for the CD26 binding CTDs is around 5 suggesting an abundance of negatively charged amino acids. However, the ACE2 binding CTDs have a theoretical pI greater than 8, suggesting a higher proportion positively charged residues. However, despite containing equivalent proportions of aromatic amino acids, the MERS-CoV CTD is significantly more hydrophobic than the others. Notably, ACE2 is localized strictly on the cell membranes, whereas DPP4 localizes on the cell membrane as well as in the cytoplasmic and extracellular fluids. The differential location of targeting receptors might be the possible reason for the lower infectivity of MERS-CoV despite having a significantly higher mortality. A comparison of the Spike Protein CTD coding sequences revealed that the SARS-CoV, MERS-CoV, and Bat-CoV had a higher GC content (~39%) than the SARS-CoV-2 (~34%). This in turn was evident from the comparison of the codon usage of the SARS-CoV-2 Spike protein wherein a significantly higher proportion of amino acids were encoded by the AT rich codons (Figure S12).

Further, we closely examined the binding regions of the four CTDs, emphasising specifically on the evolution of Y453 (Figures 5A & 5B). The residues 449-456 of the Spike Protein from SARS-CoV-2 are Asn-Tyr-Leu-Tyr-Arg-Leu-Phe-Arg. The aligned residues for this stretch from SARS-CoV are Asn-Tyr-Lys-Tyr-Arg-Tyr-Leu-Arg. The triad Tyr-Arg-Tyr in SARS-CoV generate a strong steric hindrance causing both the tyrosine residues to remain buried within the CTD (Figure 2A, middle panel). Interestingly, mutating the second tyrosine a similar yet smaller amino acid leucine (Tyr-Arg-Leu) in SARS-CoV-2 reduces the steric hindrance (Figure 2A, top panel). This allowed the otherwise buried Tyr453 to interact with amino acids from ACE2, resulting in an enhanced binding. However in MERS-CoV and Bat-CoV, these triads are present as Tyr-Ile-Asn and Tyr-Arg-Ser, respectively. This decrease in hydropathicity significantly reduces the binding affinity of MERS-CoV and Bat-CoV spike proteins with ACE2 under physiological condition. However, this in turn enables it to bind CD26 with a stronger affinity, suggesting a positive selection. Taken together; the removal of the second tyrosine from the triad to a weakly hydrophobic and smaller amino acid suggests a purifying selection in the SARS-CoV-2 Spike protein.

**Figure 5:**
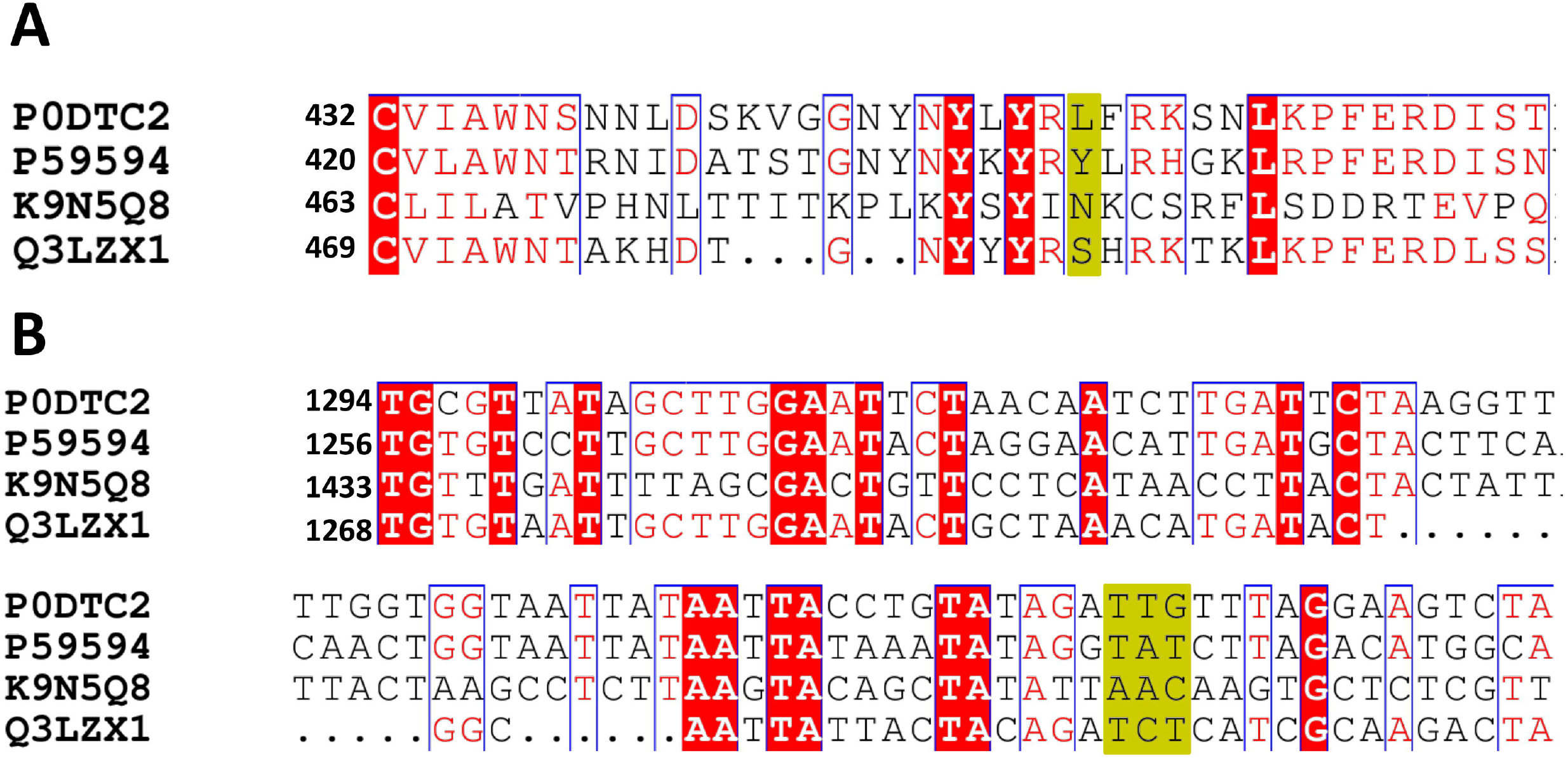
Exploring the selection pressure on the Spike protein CTDs of SARS-CoV2, SARS-CoV, MERS-CoV and Bat-CoV. **(A)** Amino acid alignment of CTDs focussing on Y453 of SARS-CoV2 spike protein and its homologous residues. **(B)** Nucleotide alignment for the ORFs coding for the CTDs..

### 4. Discussion

Although there are several reports on the evolutionary and outbreak trends of SARS-CoV-2 (Acter et al., 2020; Holmes and Rambaut, 2004; Luk et al., 2019; Shi and Wang, 2011), there are no reports so far describing the comparative molecular basis of the evolution, overtly for the spike protein CTDs with respect to the immediate and similar candidates such as Bat-CoV, SARS-CoV, and MERS-CoV. To establish and understand the molecular basis of infection for this pandemic strain along with other related coronaviruses are of utmost importance in the areas of-a) developing intervention, b) predicting the next strain in the course of evolution and c) develop an understanding of the changes in the virulence of the pandemic strains by analyzing the mutations observed. A recent report has established various mutations that are occurring in the pandemic strains and the current report can easily be extrapolated to include all the newer mutations to provide further insights into the virulence trend of the mutated strains (van Dorp et al., 2020). The novelty of the current study is that we have compared 45 verified sequences of Spike proteins from related coronaviruses to understand the unique and conserved segments of the spike proteins in order to decipher the structural and sequence similarities among the related coronaviruses to build a correlative basis.

The current study compared the spike protein sequences at amino acid (aa) and genomic nucleotide (nt) sequences. The comparison unequivocally established the corroboration of two independent analyses (aa and nt levels) and the analysis validated the results to reveal fact that SARS-CoV-2 is more similar to SARS-CoV-1 than other two closely related coronaviruses (Bat-CoV and MERS-CoV). Also, the significant conservation of amino in the S2 fragment of the Spike protein might be indicative of the conservation of the viral entry mechanism utilized by various coronaviruses irrespective of the host identity and the corresponding host receptor molecule. Our analysis also indicates a purifying selection driving the evolution of SARS-CoV-2. Previous studies have demonstrated a positive effect of the GC content on the *dN* value, thereby driving a positive selection (Du et al., 2018; Meunier and Duret, 2004). This further strengthens the theory of a purifying selection, as evident by the lower GC content in SARS-CoV-2 Spike protein coding sequence. A higher GC content exerts higher energy demands at the nucleotide level while compensating for same by coding for energy efficient amino acids. This in turn makes the replication of GC content genomes energy consuming, thus slowing the process genome replication and resultant viral replication. Structural, energetics and *in silico* mutational analyses further confirm the observation and establish the properties and locations of the binding residues Tyr453 to explain the higher affinity of SARS-CoV-2 for ACE2 receptor. We propose that the evolution of SARS-CoV-2 occurred in parallel yet independently of MERS-CoV, following a set of recombination and mutational events involving the genomes of the Bat-CoV and the SARS-CoV.

## 5. Conclusions

It is important to mention that the current study is the first of its kind to establish a comparative molecular basis on the evolution of SARS-CoV-2 to acquire its virulence and infectivity over other related coronaviruses. The work has the potential to merge with the other published data and transform the knowledgebase on SARS-CoV-2 in a newer dimension to predict upcoming outcomes and aid in the development of novel and effective interventions.

## Supplementary Materials

The supplementary data are given in a separate compiled file. A brief description of the supplementary data. Figure S1: amino acid sequence, Figure S2: A maximum parsimony phylogeny tree, Figure S3: spike protein comparison, Figure S4: Superimposition of CTDs, Figure S5: Superimposition of NTDs, Figure S6: Surface representation of protein complex, Figure S7: Comparison of maximum parsimony and maximum likelihood, Figure S8: Heatmap of hierarchical clustering, Figure S9: Heatmap of dS values, Figure S10: multiple protein sequence alignment, Figure S11: multiple nucleotide sequence alignment, Figure S12: Comparison of codon usage, Table S1: List of spike proteins.

## Author Contributions

KM and AD contributed equally to this work by performing all data analyses required. MA helped in MD simulation. PP supported the work by supervising KM; and BB supported the work by supervising AD and MA; and helped finalizing the manuscript. AD and PA drafted the manuscript. PA conceptualized, planned the execution and initiated the collaboration among NISER, NII and ILS and finalized the manuscript.

## Funding

The current work was not supported by any extramural funding.

## Acknowledgments

The infrastructural facility required to execute this study was supported by NISER, NII and ILS. KM and AD contributed equally to this work by performing all data analyses required. MA helped in MD simulation. PP supported the work by supervising KM; and BB supported the work by supervising AD and MA; and helped finalizing the manuscript. AD and PA drafted the manuscript. PA conceptualized, planned the execution and initiated the collaboration among NISER, NII and ILS and finalized the manuscript.

## Conflicts of Interest

Authors declare no conflict of interest.

## Notes

### Competing Interest Statement

The authors have declared no competing interest.

## References

Acter, T., Uddin, N., Das, J., Akhter, A., Choudhury, T.R., Kim, S., 2020. Evolution of severe acute respiratory syndrome coronavirus 2 (SARS-CoV-2) as coronavirus disease 2019 (COVID-19) pandemic: A global health emergency. Sci. Total Environ. https://doi.org/10.1016/j.scitotenv.2020.138996

Andersen, K.G., Rambaut, A., Lipkin, W.I., Holmes, E.C., Garry, R.F., 2020. The proximal origin of SARS-CoV-2. Nat. Med. https://doi.org/10.1038/s41591-020-0820-9

Ashour, H.M., Elkhatib, W.F., Rahman, M.M., Elshabrawy, H.A., 2020. Insights into the recent 2019 novel coronavirus (Sars-coV-2) in light of past human coronavirus outbreaks. Pathogens. https://doi.org/10.3390/pathogens9030186

Baker, F.B., 1974. Stability of two hierarchical grouping techniques case I: Sensitivity to data errors. J. Am. Stat. Assoc. 69, 440–445. https://doi.org/10.1080/01621459.1974.10482971

Baskaran, K., Duarte, J.M., Biyani, N., Bliven, S., Capitani, G., 2014. A PDB-wide, evolution-based assessment of protein-protein interfaces. BMC Struct. Biol. 14. https://doi.org/10.1186/s12900-014-0022-0

Benvenuto, D., Giovanetti, M., Ciccozzi, A., Spoto, S., Angeletti, S., Ciccozzi, M., 2020. The 2019-new coronavirus epidemic: Evidence for virus evolution. J. Med. Virol. 92, 455–459. https://doi.org/10.1002/jmv.25688

Charif, D., Lobry, J.R., 2007. SeqinR 1.0-2: A Contributed Package to the R Project for Statistical Computing Devoted to Biological Sequences Retrieval and Analysis. pp. 207–232. https://doi.org/10.1007/978-3-540-35306-5_10

DeLano, W.L., 2020. The PyMOL Molecular Graphics System, Version 2.3. Schrödinger LLC. https://doi.org/10.1038/hr.2014.17

Du, M.Z., Zhang, C., Wang, H., Liu, S., Wei, W., Guo, F.B., 2018. The GC content as a main factor shaping the amino acid usage during bacterial evolution process. Front. Microbiol. 9. https://doi.org/10.3389/fmicb.2018.02948

Edgar, R.C., 2004. MUSCLE: Multiple sequence alignment with high accuracy and high throughput. Nucleic Acids Res. 32, 1792–1797. https://doi.org/10.1093/nar/gkh340

Galili, T., 2015. dendextend: An R package for visualizing, adjusting and comparing trees of hierarchical clustering. Bioinformatics 31, 3718–3720. https://doi.org/10.1093/bioinformatics/btv428

Giribet, G., 2005. TNT: Tree Analysis Using New Technology. Syst. Biol. 54, 176–178. https://doi.org/10.1080/10635150590905830

Gouet, P., Robert, X., Courcelle, E., 2005. ESPript/ENDscript: sequence and 3D information from protein structures. Acta Crystallogr. Sect. A Found. Crystallogr. 61, c42–c43. https://doi.org/10.1107/s0108767305098211

Heo, L., Lee, H., Seok, C., 2016. GalaxyRefineComplex: Refinement of protein-protein complex model structures driven by interface repacking. Sci. Rep. 6. https://doi.org/10.1038/srep32153

Holmes, E.C., Rambaut, A., 2004. Viral evolution and the emergence of SARS coronavirus. Philos. Trans. R. Soc. London. Ser. B Biol. Sci. 359, 1059–1065. https://doi.org/10.1098/rstb.2004.1478

Ingram, T., Mahler, D.L., 2013. SURFACE: Detecting convergent evolution from comparative data by fitting Ornstein-Uhlenbeck models with stepwise Akaike Information Criterion. Methods Ecol. Evol. 4, 416–425. https://doi.org/10.1111/2041-210X.12034

Jin, Y., Yang, H., Ji, W., Wu, W., Chen, S., Zhang, W., Duan, G., 2020. Virology, epidemiology, pathogenesis, and control of covid-19. Viruses. https://doi.org/10.3390/v12040372

Krüger, D.M., Gohlke, H., 2010. DrugScorePPI webserver: Fast and accurate in silico alanine scanning for scoring protein-protein interactions. Nucleic Acids Res. 38. https://doi.org/10.1093/nar/gkq471

Kumar, S., Stecher, G., Li, M., Knyaz, C., Tamura, K., 2018. MEGA X: Molecular evolutionary genetics analysis across computing platforms. Mol. Biol. Evol. 35, 1547–1549. https://doi.org/10.1093/molbev/msy096

Lapointe, F.J., Legendre, P., 1995. Comparison tests for dendrograms: A comparative evaluation. J. Classif. https://doi.org/10.1007/BF03040858

Li, F., 2016. Structure, Function, and Evolution of Coronavirus Spike Proteins. Annu. Rev. Virol. 3, 237–261. https://doi.org/10.1146/annurev-virology-110615-042301

Li, Z., Natarajan, P., Ye, Y., Hrabe, T., Godzik, A., 2014. POSA: A user-driven, interactive multiple protein structure alignment server. Nucleic Acids Res. 42. https://doi.org/10.1093/nar/gku394

Luk, H.K.H., Li, X., Fung, J., Lau, S.K.P., Woo, P.C.Y., 2019. Molecular epidemiology, evolution and phylogeny of SARS coronavirus. Infect. Genet. Evol. https://doi.org/10.1016/j.meegid.2019.03.001

Meunier, J., Duret, L., 2004. Recombination drives the evolution of GC-content in the human genome. Mol. Biol. Evol. 21, 984–990. https://doi.org/10.1093/molbev/msh070

Moreira, I.S., Koukos, P.I., Melo, R., Almeida, J.G., Preto, A.J., Schaarschmidt, J., Trellet, M., Gümüş, Z.H., Costa, J., Bonvin, A.M.J.J., 2017. SpotOn: High Accuracy Identification of Protein-Protein Interface Hot-Spots. Sci. Rep. 7. https://doi.org/10.1038/s41598-017-08321-2

Negi, S.S., Schein, C.H., Oezguen, N., Power, T.D., Braun, W., 2007. InterProSurf: A web server for predicting interacting sites on protein surfaces. Bioinformatics 23, 3397–3399. https://doi.org/10.1093/bioinformatics/btm474

Pierce, B.G., Wiehe, K., Hwang, H., Kim, B.H., Vreven, T., Weng, Z., 2014. ZDOCK server: Interactive docking prediction of protein-protein complexes and symmetric multimers. Bioinformatics 30, 1771–1773. https://doi.org/10.1093/bioinformatics/btu097

Posada, D., Crandall, K.A., 1998. MODELTEST: Testing the model of DNA substitution. Bioinformatics 14, 817–818. https://doi.org/10.1093/bioinformatics/14.9.817

Rocha, E.P.C., Smith, J.M., Hurst, L.D., Holden, M.T.G., Cooper, J.E., Smith, N.H., Feil, E.J., 2006. Comparisons of dN/dS are time dependent for closely related bacterial genomes. J. Theor. Biol. 239, 226–235. https://doi.org/10.1016/j.jtbi.2005.08.037

Rodrigues, C.H.M., Myung, Y., Pires, D.E.V., Ascher, D.B., 2019. MCSM-PPI2: predicting the effects of mutations on protein-protein interactions. Nucleic Acids Res. 47, W338–W344. https://doi.org/10.1093/nar/gkz383

Rothan, H.A., Byrareddy, S.N., 2020. The epidemiology and pathogenesis of coronavirus disease (COVID-19) outbreak. J. Autoimmun. https://doi.org/10.1016/j.jaut.2020.102433

Schliep, K.P., 2011. phangorn: Phylogenetic analysis in R. Bioinformatics 27, 592–593. https://doi.org/10.1093/bioinformatics/btq706

Shi, Z., Wang, L.F., 2011. Evolution of SARS Coronavirus and the relevance of modern Molecular Epidemiology, in: Genetics and Evolution of Infectious Diseases. Elsevier Inc., pp. 711–728. https://doi.org/10.1016/B978-0-12-384890-1.00027-3

Sievers, F., Higgins, D.G., 2014. Clustal Omega. Curr. Protoc. Bioinforma. 2014, 3.13.1-3.13.16. https://doi.org/10.1002/0471250953.bi0313s48

Smith, M.R., 2020. Information theoretic Generalized Robinson-Foulds metrics for comparing phylogenetic trees. Bioinformatics. https://doi.org/10.1093/bioinformatics/btaa614

Sun, J., He, W.T., Wang, L., Lai, A., Ji, X., Zhai, X., Li, G., Suchard, M.A., Tian, J., Zhou, J., Veit, M., Su, S., 2020. COVID-19: Epidemiology, Evolution, and Cross-Disciplinary Perspectives. Trends Mol. Med. 26, 483–495. https://doi.org/10.1016/j.molmed.2020.02.008

van Dorp, L., Acman, M., Richard, D., Shaw, L.P., Ford, C.E., Ormond, L., Owen, C.J., Pang, J., Tan, C.C.S., Boshier, F.A.T., Ortiz, A.T., Balloux, F., 2020. Emergence of genomic diversity and recurrent mutations in SARS-CoV-2. Infect. Genet. Evol. 83, 104351. https://doi.org/10.1016/j.meegid.2020.104351

Veeramalai, M., Ye, Y., Godzik, A., 2008. TOPS++FATCAT: Fast flexible structural alignment using constraints derived from TOPS+ Strings Model. BMC Bioinformatics 9. https://doi.org/10.1186/1471-2105-9-358

Walls, A.C., Park, Y.-J., Tortorici, M.A., Wall, A., McGuire, A.T., Veesler, D., 2020. Structure, Function, and Antigenicity of the SARS-CoV-2 Spike Glycoprotein. Cell. https://doi.org/10.1016/j.cell.2020.02.058

Wang, Qihui, Zhang, Y., Wu, L., Niu, S., Song, C., Zhang, Z., Lu, G., Qiao, C., Hu, Y., Yuen, K.Y., Wang, Qisheng, Zhou, H., Yan, J., Qi, J., 2020. Structural and Functional Basis of SARS-CoV-2 Entry by Using Human ACE2. Cell 181, 894–904.e9. https://doi.org/10.1016/j.cell.2020.03.045

Wernersson, R., Pedersen, A.G., 2003. RevTrans: Multiple alignment of coding DNA from aligned amino acid sequences. Nucleic Acids Res. 31, 3537–3539. https://doi.org/10.1093/nar/gkg609

WHO, 2020. Coronavirus disease 2019 (COVID-19) Situation Report – 174.

Wrapp, D., Wang, N., Corbett, K.S., Goldsmith, J.A., Hsieh, C.-L., Abiona, O., Graham, B.S., McLellan, J.S., 2020. Cryo-EM structure of the 2019-nCoV spike in the prefusion conformation. Science (80-.). 367, 1260LP–1263. https://doi.org/10.1126/science.abb2507

Xue, L.C., Rodrigues, J.P., Kastritis, P.L., Bonvin, A.M., Vangone, A., 2016. PRODIGY: A web server for predicting the binding affinity of protein-protein complexes. Bioinformatics 32, 3676–3678. https://doi.org/10.1093/bioinformatics/btw514

Yan, R., Zhang, Y., Li, Y., Xia, L., Guo, Y., Zhou, Q., 2020. Structural basis for the recognition of SARS-CoV-2 by full-length human ACE2. Science (80-.). 367, 1444–1448. https://doi.org/10.1126/science.abb2762

Yu, G., Smith, D.K., Zhu, H., Guan, Y., Lam, T.T.Y., 2017. Ggtree: an R Package for Visualization and Annotation of Phylogenetic Trees With Their Covariates and Other Associated Data. Methods Ecol. Evol. 8, 28–36. https://doi.org/10.1111/2041-210X.12628

Zhao, J., Yang, Y., Huang, H.-P., Li, D., Gu, D.-F., Lu, X.-F., Zhang, Z., Liu, L., Liu, T., Liu, Y.-K., He, Y.-J., Sun, B., Wei, M.-L., Yang, G.-Y., Wang, X., Zhang, L., Zhou, X.-Y., Xing, M.-Z., Wang, P.G., 2020. Relationship between the ABO Blood Group and the COVID-19 Susceptibility. medRxiv 2020.03.11.20031096. https://doi.org/10.1101/2020.03.11.20031096

