## Supplemental data for "Molecular basis of the logical evolution of the novel coronavirus SARS-CoV-2: A comparative analysis"

***Running title:*** **molecular basis of SARS-CoV-2 emergence**

### SUPPLEMENTARY INFORMATION

### Figures, Figure Legends and Tables

Figure S1

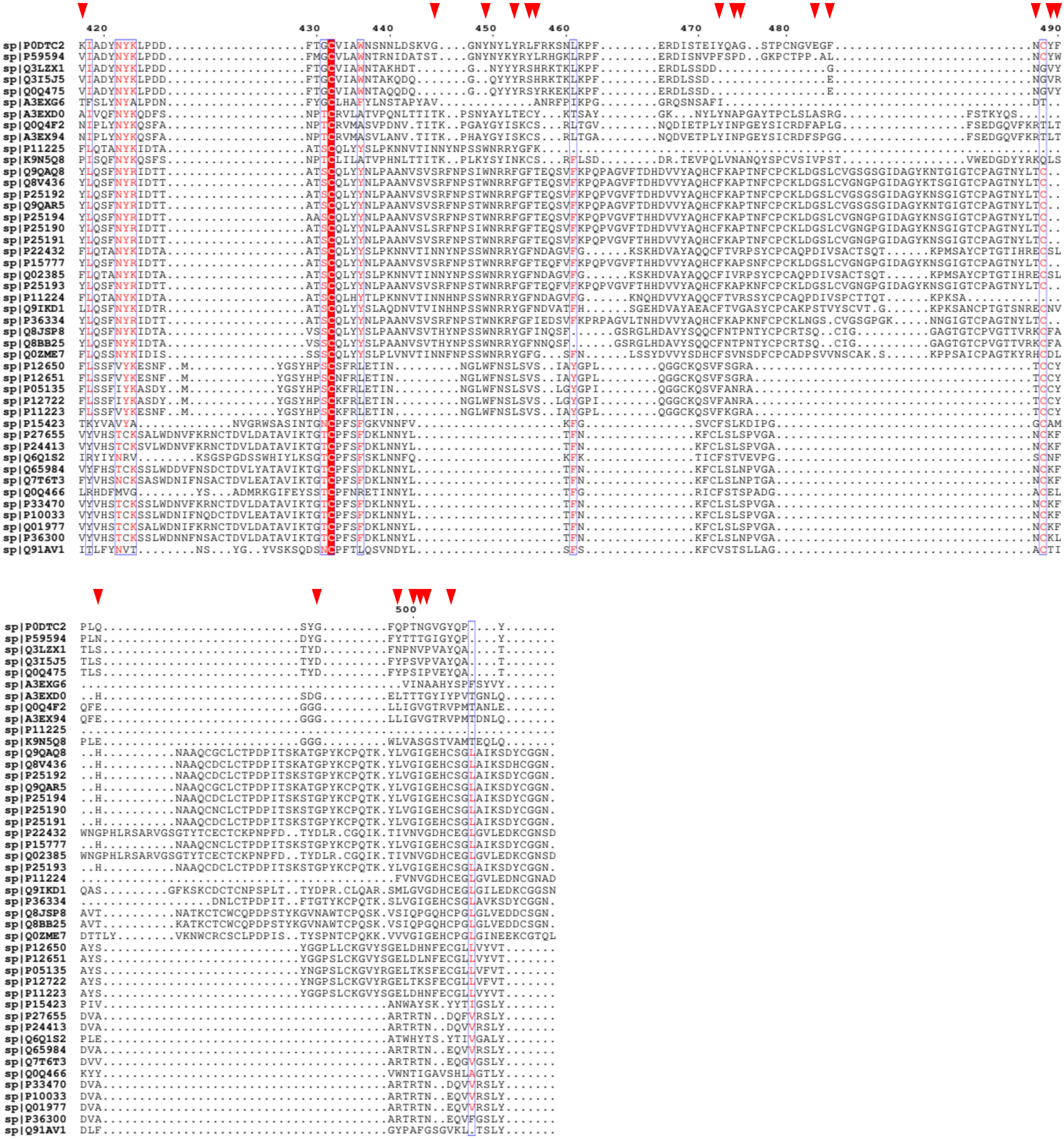
420
430
440
450
460
470
480
490
500
sp|P0DTC2
sp|P59594
sp|Q3LZX1
sp|Q3I5J5
sp|Q0Q475
sp|A3EXG6
sp|A3EXD0
sp|Q0Q4F2
sp|A3EX94
sp|P11225
sp|K9N5Q8
sp|Q9QAQ8
sp|Q8V436
sp|P25192
sp|Q9QAR5
sp|P25194
sp|P25190
sp|P25191
sp|P22432
sp|P15777
sp|Q02385
sp|P25193
sp|P11224
sp|Q9IKD1
sp|P36334
sp|Q8JSP8
sp|Q8BB25
sp|Q0ZME7
sp|P12650
sp|P12651
sp|P05135
sp|P12722
sp|P11223
sp|P15423
sp|P27655
sp|P24413
sp|Q6Q1S2
sp|Q65984
sp|Q7T6T3
sp|Q0Q466
sp|P33470
sp|P10033
sp|Q01977
sp|P36300
sp|Q91AV1

**FIGURE S1:** A comparison of the amino acids sequences of Spike proteins from various coronaviruses (the figure spans the ACE2 binding region of the protein from SARS-CoV2 only). The first sequence (P0DTC2) is that of the Spike protein from SARS-CoV2. The residues of Spike protein from SARS-CoV2 that interacts with human ACE2 are marked with red triangles.

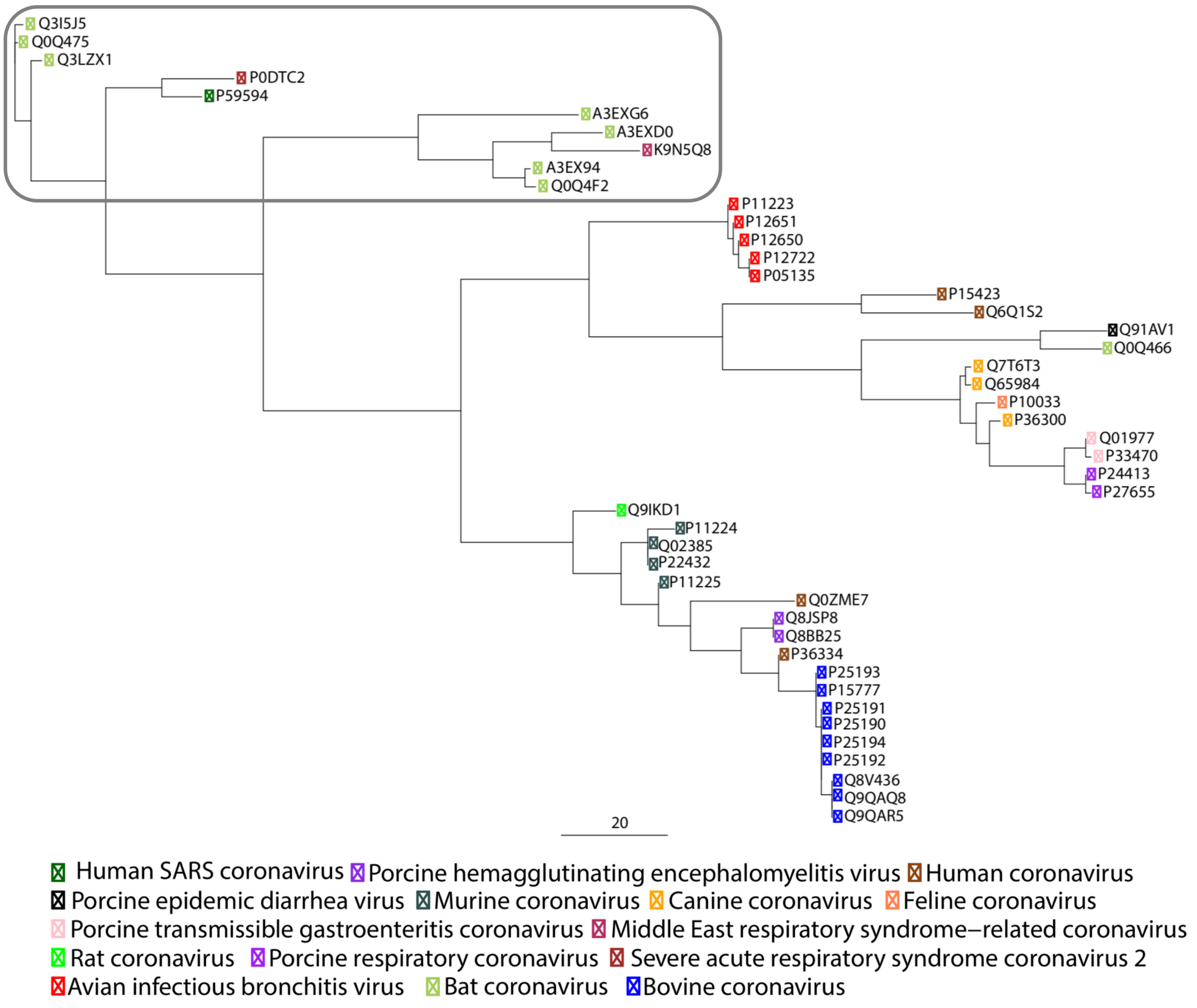
Q3I5J5
Q0Q475
Q3LZX1
P0DTC2
P59594
A3EXG6
A3EXD0
K9N5Q8
A3EX94
Q0Q4F2
P11223
P12651
P12650
P12722
P05135
P15423
Q6Q1S2
Q91AV1
Q0Q466
Q7T6T3
Q65984
P10033
P36300
Q01977
P33470
P24413
P27655
Q9IKD1
P11224
Q02385
P22432
P11225
Q0ZME7
Q8JSP8
Q8BB25
P36334
P25193
P15777
P25191
P25190
P25194
P25192
Q8V436
Q9QAQ8
Q9QAR5
20
Human SARS coronavirus
Porcine hemagglutinating encephalomyelitis virus
Human coronavirus
Porcine epidemic diarrhea virus
Murine coronavirus
Canine coronavirus
Feline coronavirus
Porcine transmissible gastroenteritis coronavirus
Middle East respiratory syndrome–related coronavirus
Rat coronavirus
Porcine respiratory coronavirus
Severe acute respiratory syndrome coronavirus 2
Avian infectious bronchitis virus
Bat coronavirus
Bovine coronavirus

**FIGURE S2:** A maximum parsimony phylogenetic tree for the Spike proteins from various coronaviruses. Human coronavirus species are underlined in red, while bat coronaviruses are underlined in blue. It is worth noticing that all human coronaviruses with significantly high morbidities (SARS-CoV (P59594), MERS-CoV (K9N5Q8) and SARS-CoV2 (P0DTC2) cluster separately along with several Bat-CoVs).

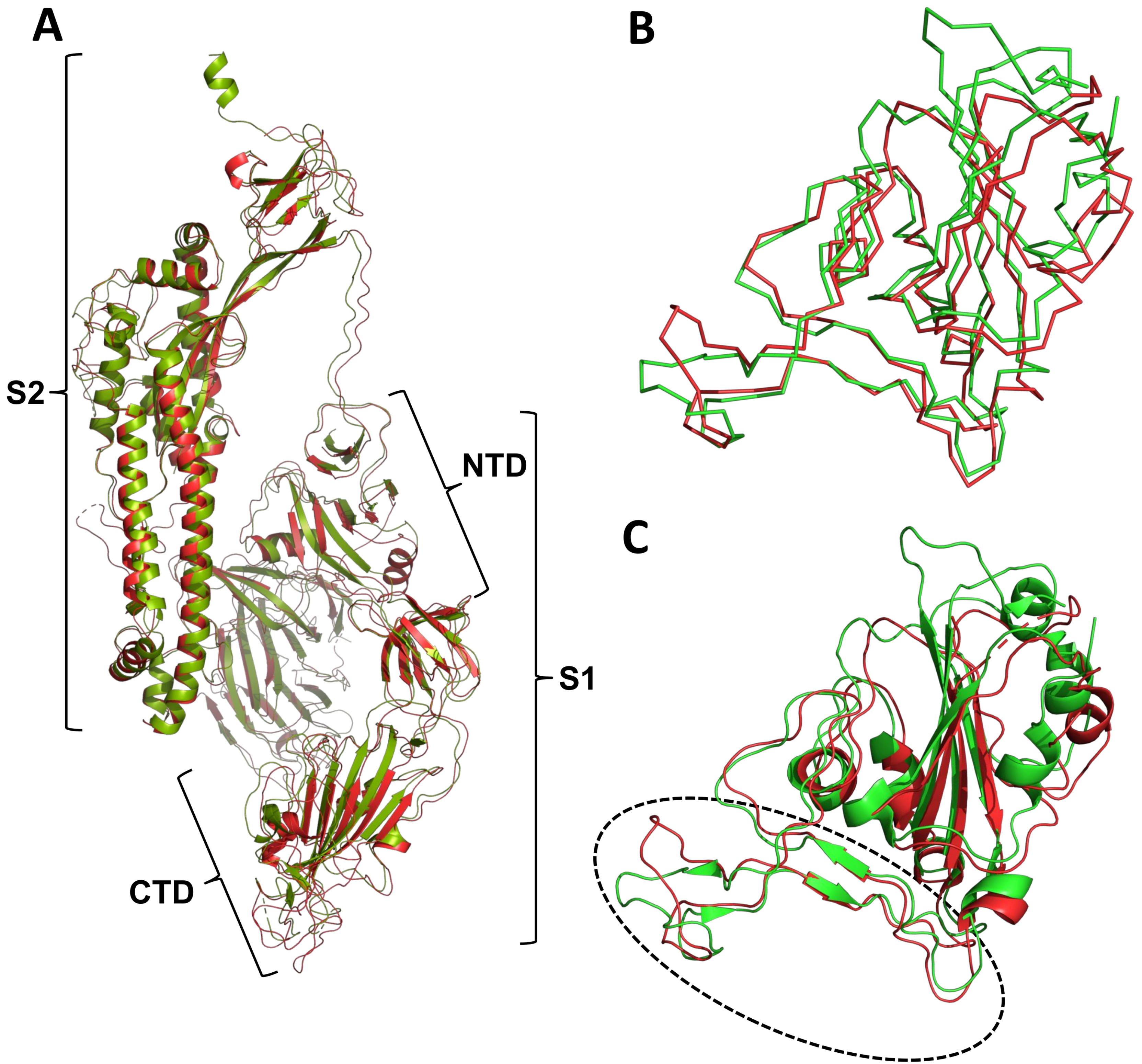
A
S2
NTD
S1
CTD
B
C

**FIGURE S3: (A)** A comparison of the Spike protein structures from SARS-CoV and SARS-CoV2 and secondary structures (right panel) (CoV2 in lime-green, CoV in tv-red). While the overall structures are globally similar, there are local yet conspicuous variations, especially in the S1 fragment region. While the CTD domain of CoV2 displays lesser number of flexible loops like structures and longer β-strands than CoV, the NTD fold remains largely. **(B & C)** A comparison of the ACE2 recognizing CTD form the fragment S1 of the Spike proteins from SARS-CoV and SARS-CoV2 – Cα chain **(B)** and secondary structures **(C)** (CoV2 in lime-green, CoV in tv-red). A significant section of the CTD from CoV2 is comprised of well-defined secondary structures with longer β-strands as compared to that of the CTD from CoV (encircled in black). It is worth mentioning that few amino acid residues from this region are in involved in recognising and binding the ACE2.

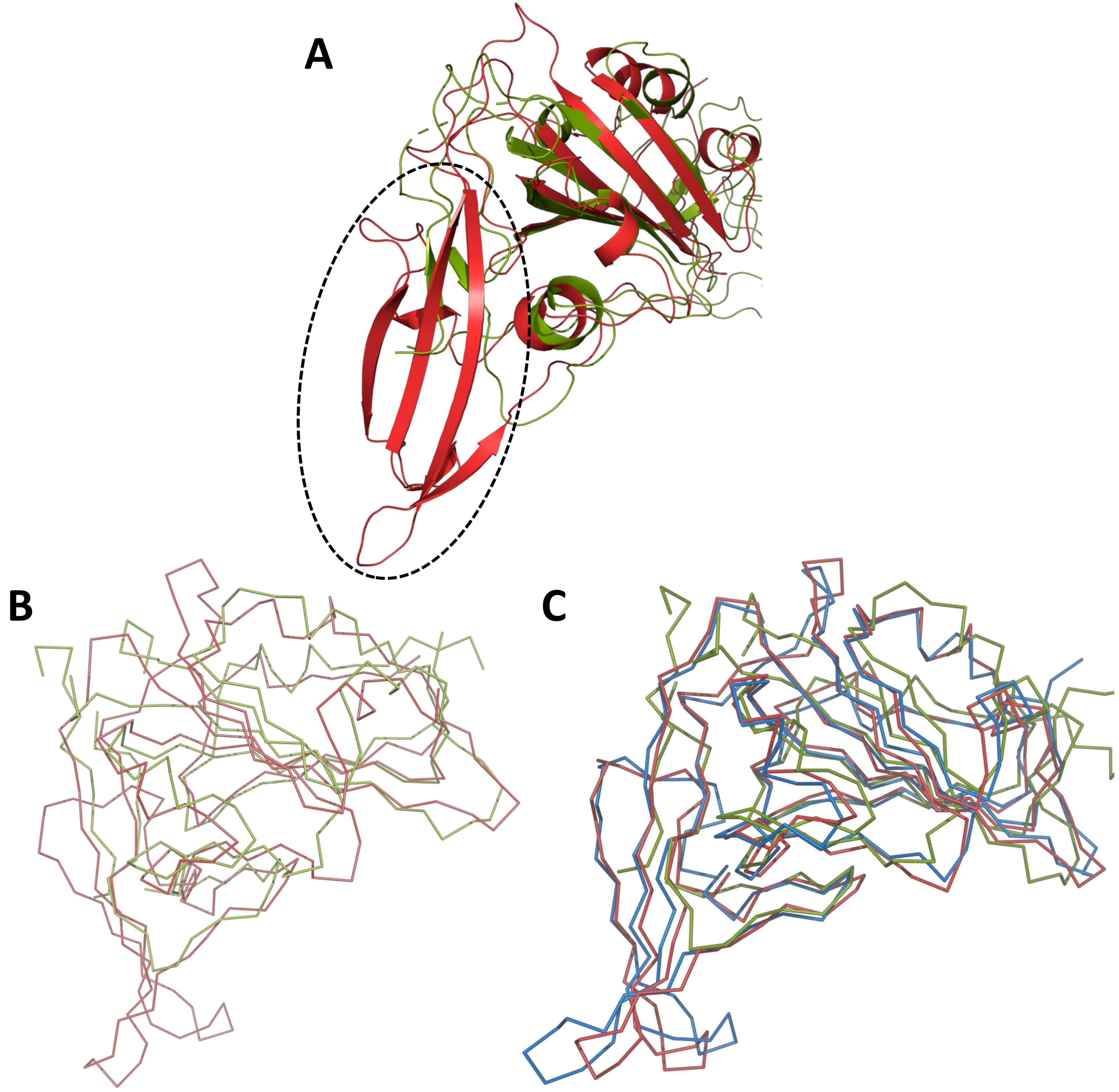
A
B
C

**FIGURE S4: (A)** A zoomed view of the superimposition of the CD26 recognizing CTD from Bat-CoV Spike protein onto the ACE2 recognizing CTD from SARS-CoV2 spike protein (SARS-CoV2 in lime-green, Bat-CoV in tv-red). The CD26 of Bat-CoV recognized domain is significant larger as compared to the ACE2 binding domain of SARS-CoV2 with and anti-parallel β-sheet protruding out. Noticeably, this β-sheet includes a number of key residues involved in recognition and binding of CD26. **(B)** A superimposition of the Cα chain of the CD26 recognizing CTD from MERS-CoV Spike protein and the ACE2 binding domain the SARS-CoV2 spike protein (SARS-CoV2 in lime-green , METS-CoV in tv-red). **(C)** A superimposition of the Cα chain of the CD26 recognizing CTD from MERS-CoV and Bat-CoV Spike proteins and the ACE2 binding domain the SARS-CoV2 spike protein (SARS-CoV2 in lime-green , METS-CoV in tv-red, Bat-CoV in tv-blue). The structural similarity of the CTD of the spike proteins from MERS-CoV and Bat-CoV is striking.

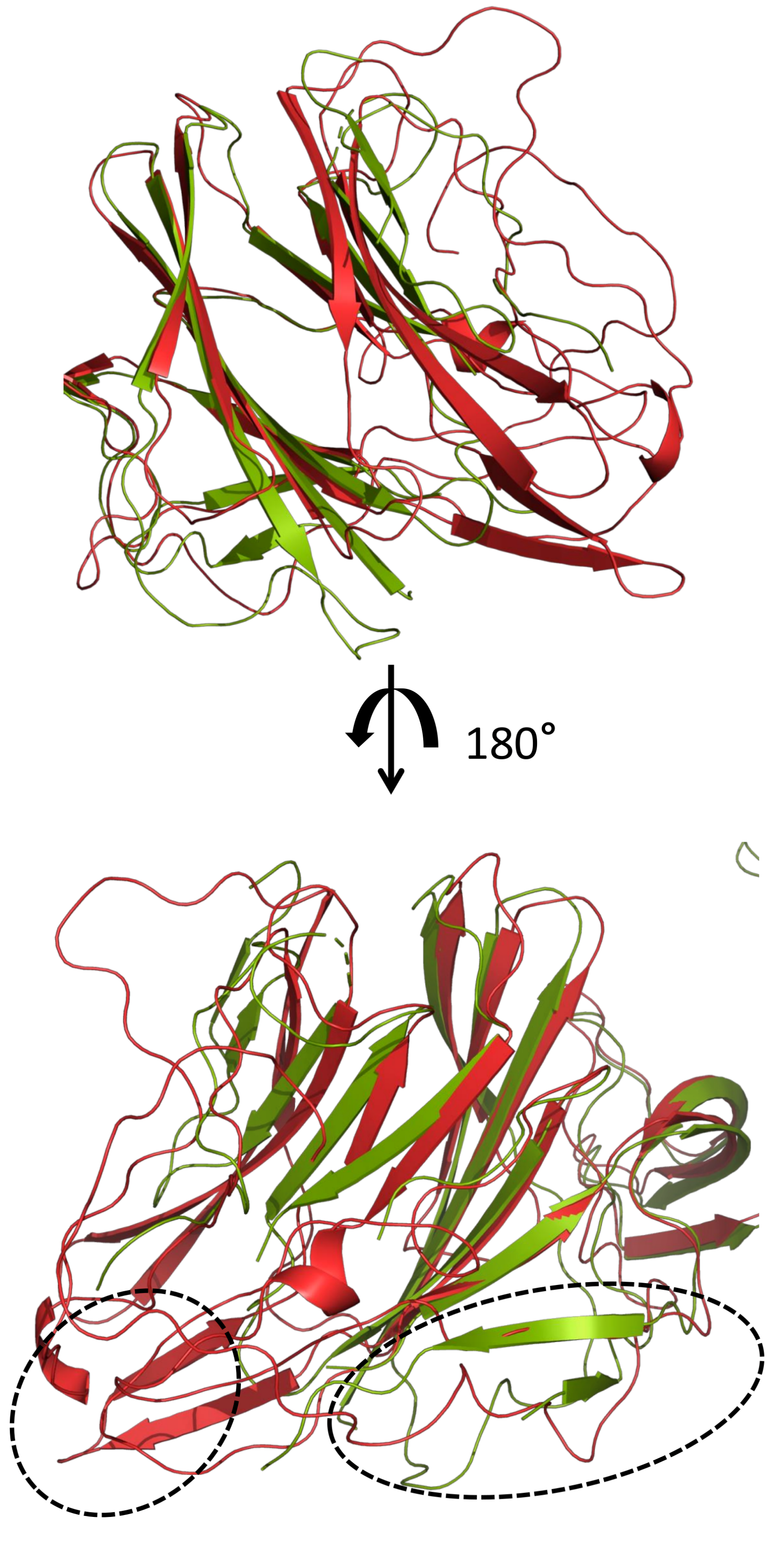
180°

**FIGURE S5:** A zoomed view superimposition of the lectin recognizing NTD from Bat-CoV Spike protein onto the NTD from SARS-CoV2 spike protein. A 180° horizontal rotation views (SARS-CoV2 in lime-green, Bov-CoV in tv-red). The Bov-CoV NTD comprises of longer β-strands, with 2 additional helices. Of Not the position of two β-strands considerably vary between the two structures, making the CoV2 NTD more compact while the Bov-CoV NTD is more stretched.

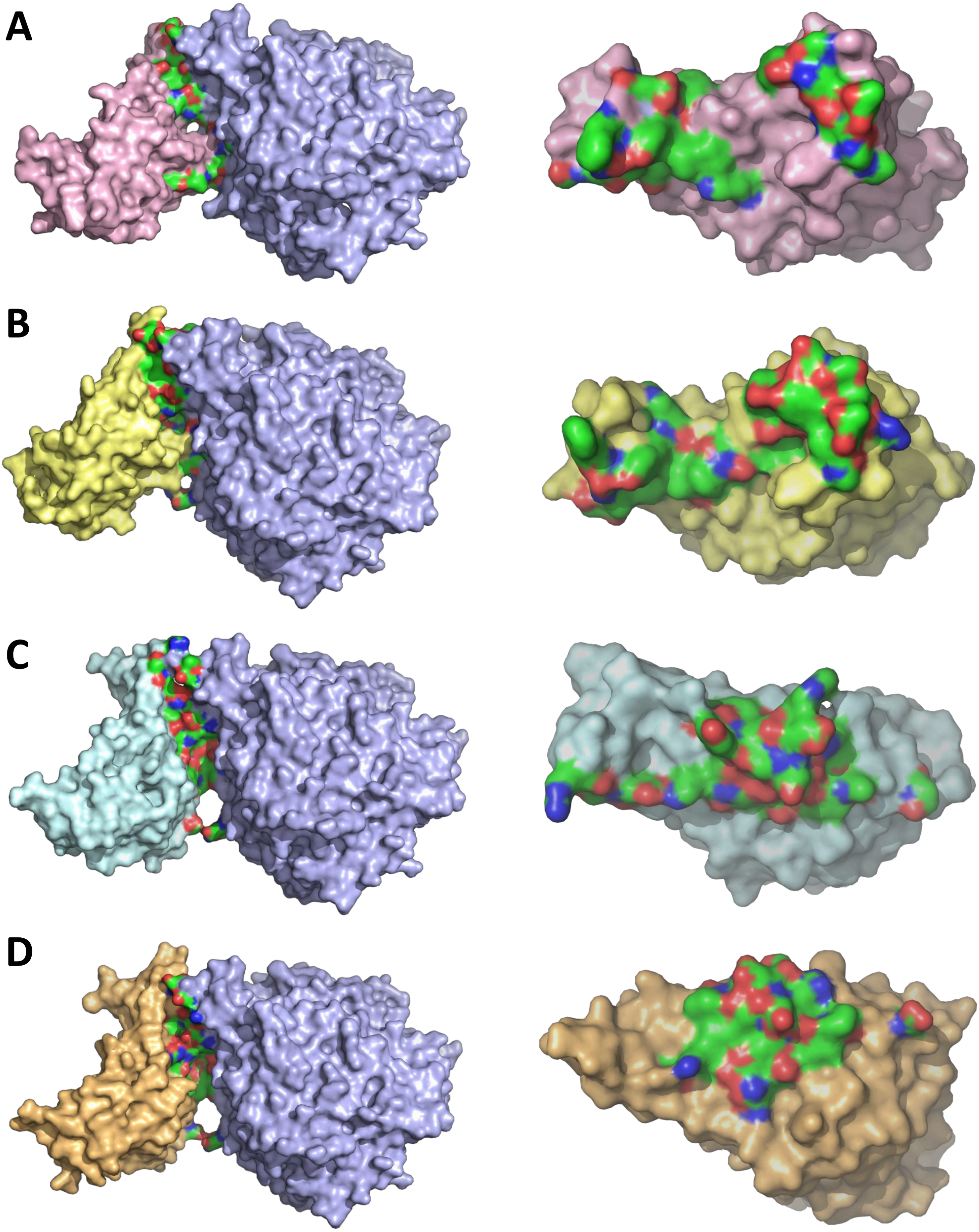

A
B
C
D

**Figure S6: Surface representation protein complex for ACE2 (ashy grey) with the CTDs** of **(A)** SARS-CoV (salmon); **(B)** SARS-CoV (tv-yellow); **(C)** MERS-CoV (light-cyan); and **(D)** Bat-CoV (light-brown). Surfaces of the interacting residues are represented in red (postively charged), blue (negatively charged) and green (uncharged) colors. The interacting face each CTD represented in the respective right panels. The distribution of interacting residues greatly varies from SARS-CoV2 on end of the spectrum to Bat-CoV at the other.

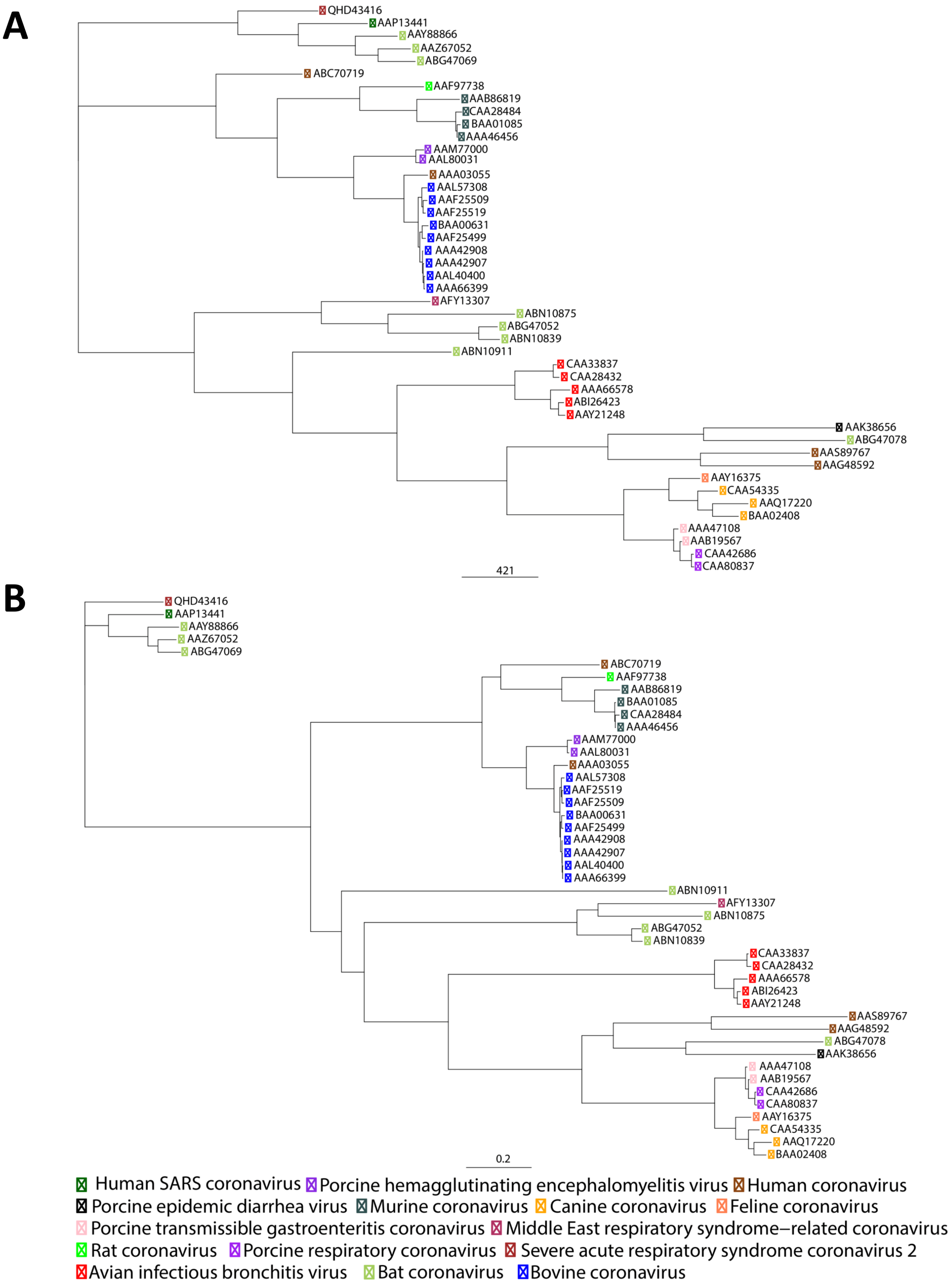
A
QHD43416
AAP13441
AAY88866
AAZ67052
ABG47069
ABC70719
AAF97738
AAB86819
CAA28484
BAA01085
AAA46456
AAM77000
AAL80031
AAA03055
AAL57308
AAF25509
AAF25519
BAA00631
AAF25499
AAA42908
AAA42907
AAL40400
AAA66399
AFY13307
ABN10875
ABG47052
ABN10839
ABN10911
CAA33837
CAA28432
AAA66578
ABI26423
AAY21248
AAK38656
ABG47078
AAS89767
AAG48592
AAY16375
CAA54335
AAQ17220
BAA02408
AAA47108
AAB19567
CAA42686
CAA80837
421
B
QHD43416
AAP13441
AAY88866
AAZ67052
ABG47069
ABC70719
AAF97738
AAB86819
BAA01085
CAA28484
AAA46456
AAM77000
AAL80031
AAA03055
AAL57308
AAF25519
AAF25509
BAA00631
AAF25499
AAA42908
AAA42907
AAL40400
AAA66399
ABN10911
AFY13307
ABN10875
ABG47052
ABN10839
CAA33837
CAA28432
AAA66578
ABI26423
AAY21248
AAS89767
AAG48592
ABG47078
AAK38656
AAA47108
AAB19567
CAA42686
CAA80837
AAY16375
CAA54335
AAQ17220
BAA02408
0.2
Human SARS coronavirus
Porcine hemagglutinating encephalomyelitis virus
Human coronavirus
Porcine epidemic diarrhea virus
Murine coronavirus
Canine coronavirus
Feline coronavirus
Porcine transmissible gastroenteritis coronavirus
Middle East respiratory syndrome−related coronavirus
Rat coronavirus
Porcine respiratory coronavirus
Severe acute respiratory syndrome coronavirus 2
Avian infectious bronchitis virus
Bat coronavirus
Bovine coronavirus

**FIGURE S7:** A comparative **(A)** maximum parsimony and **(B)** maximum likelihood phylogenetic analysis of Spike proteins.

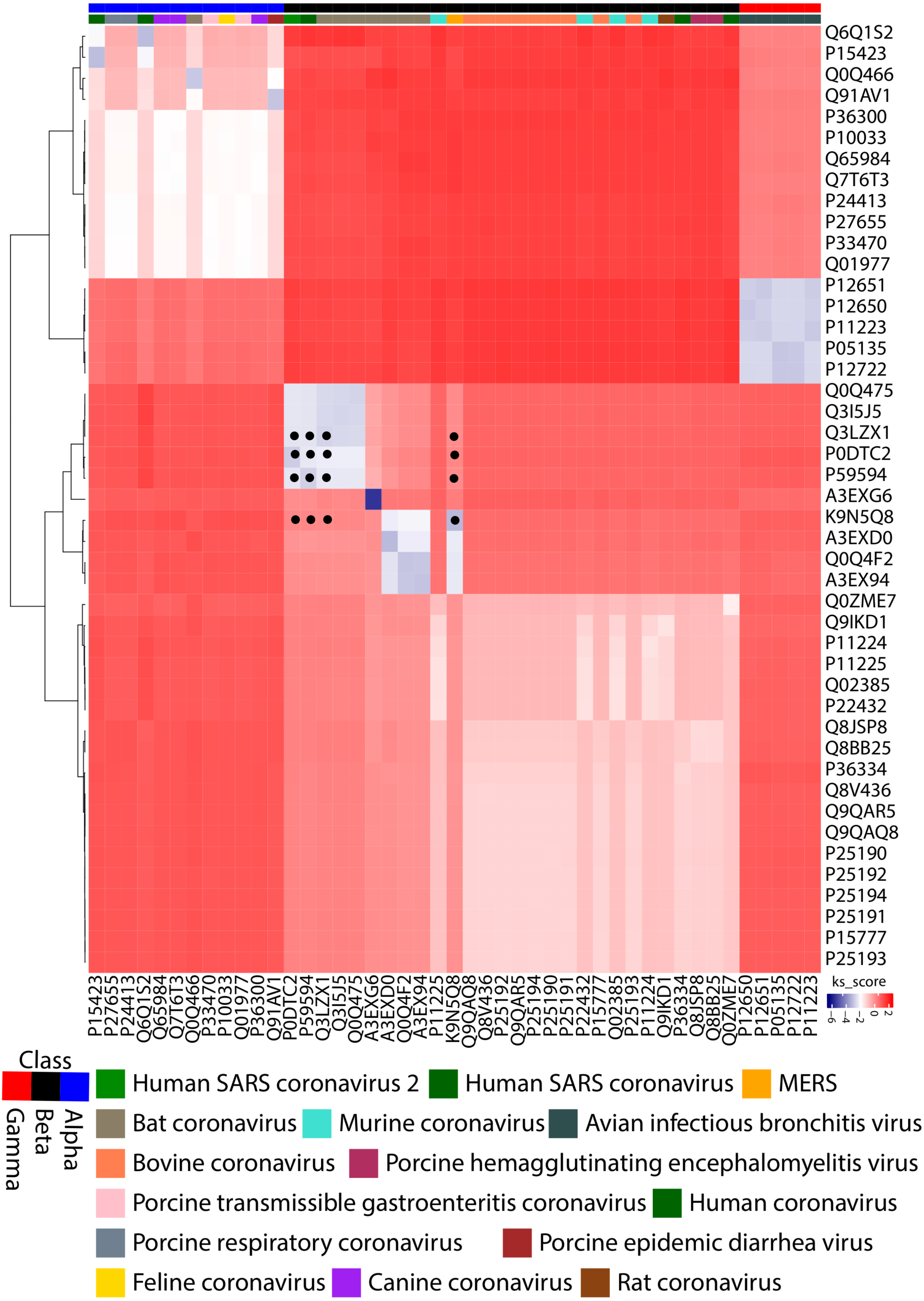
Q6Q1S2
P15423
Q0Q466
Q91AV1
P36300
P10033
Q65984
Q7T6T3
P24413
P27655
P33470
Q01977
P12651
P12650
P11223
P05135
P12722
Q0Q475
Q3I5J5
Q3LZX1
P0DTC2
P59594
A3EXG6
K9N5Q8
A3EXD0
Q0Q4F2
A3EX94
Q0ZME7
Q9IKD1
P11224
P11225
Q02385
P22432
Q8JSP8
Q8BB25
P36334
Q8V436
Q9QAR5
Q9QAQ8
P25190
P25192
P25194
P25191
P15777
P25193
P15423
P27655
P24413
Q6Q1S2
Q65984
Q7T6T3
Q0Q466
P33470
P10033
Q01977
P36300
Q91AV1
P0DTC2
P59594
Q3LZX1
Q3I5J5
Q0Q475
A3EXG6
A3EXD0
Q0Q4F2
A3EX94
P11225
K9N5Q8
Q9QAQ8
Q8V436
P25192
Q9QAR5
P25194
P25190
P25191
P22432
P15777
Q02385
P25193
P11224
Q9IKD1
P36334
Q8JSP8
Q8BB25
Q0ZME7
P12650
P12651
P05135
P12722
P11223
ks_score
−6
−4
−2
0
2
Class
Alpha
Beta
Gamma
Human SARS coronavirus 2
Human SARS coronavirus
MERS
Bat coronavirus
Murine coronavirus
Avian infectious bronchitis virus
Bovine coronavirus
Porcine hemagglutinating encephalomyelitis virus
Porcine transmissible gastroenteritis coronavirus
Human coronavirus
Porcine respiratory coronavirus
Porcine epidemic diarrhea virus
Feline coronavirus
Canine coronavirus
Rat coronavirus

**FIGURE S8:** A heat map depicting the hierarchal clustering of the *dS* values of the Spike protein encoding ORFs (Values within SARS-CoV2, SARS-CoV, MERS-CoV and Bat-CoV are marked with a black dot).

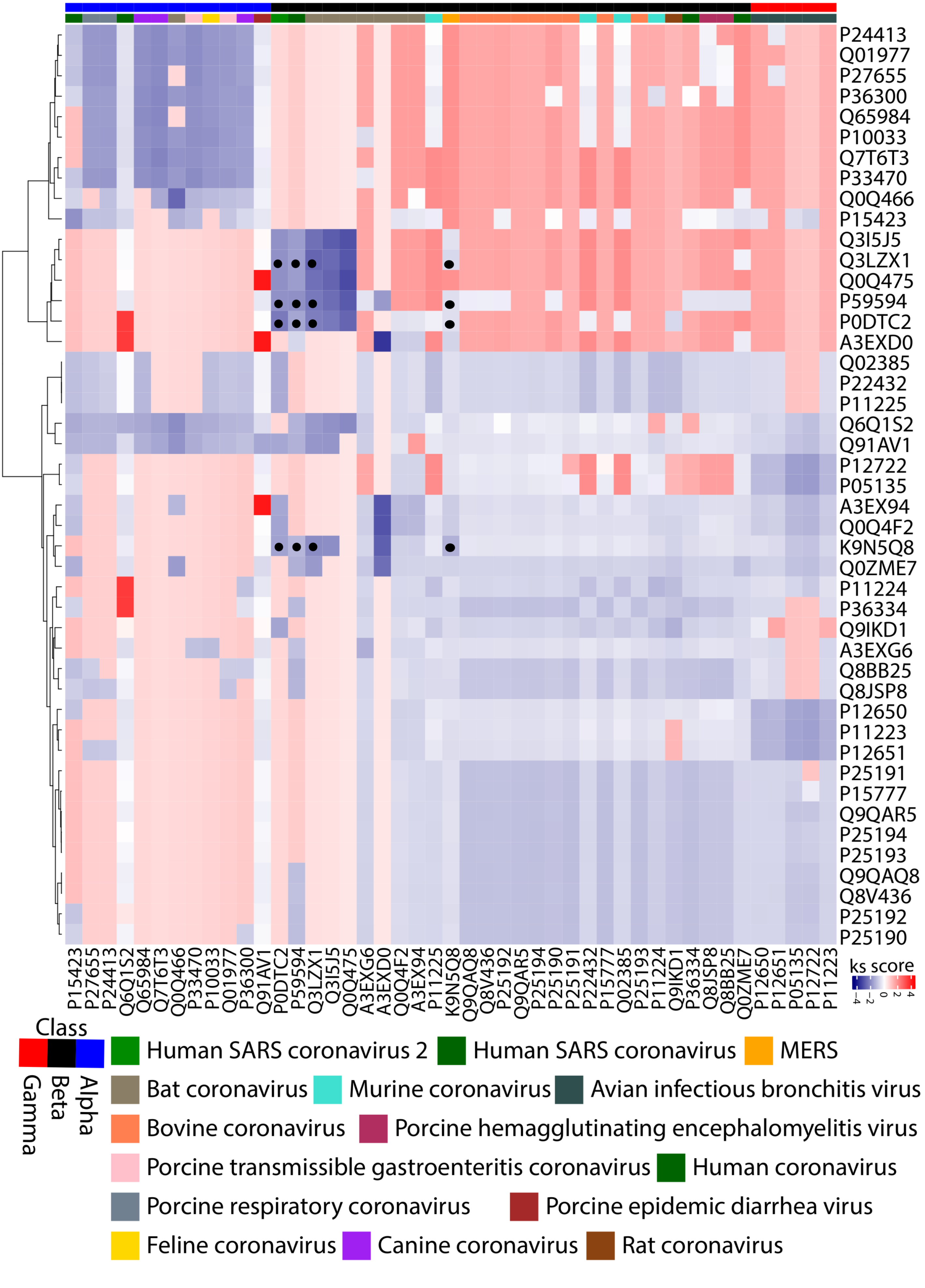

P24413
Q01977
P27655
P36300
Q65984
P10033
Q7T6T3
P33470
Q0Q466
P15423
Q3I5J5
Q3LZX1
Q0Q475
P59594
P0DTC2
A3EXD0
Q02385
P22432
P11225
Q6Q1S2
Q91AV1
P12722
P05135
A3EX94
Q0Q4F2
K9N5Q8
Q0ZME7
P11224
P36334
Q9IKD1
A3EXG6
Q8BB25
Q8JSP8
P12650
P11223
P12651
P25191
P15777
Q9QAR5
P25194
P25193
Q9QAQ8
Q8V436
P25192
P25190
ks score
Class
Alpha
Beta
Gamma
Human SARS coronavirus 2
Human SARS coronavirus
MERS
Bat coronavirus
Murine coronavirus
Avian infectious bronchitis virus
Bovine coronavirus
Porcine hemagglutinating encephalomyelitis virus
Porcine transmissible gastroenteritis coronavirus
Human coronavirus
Porcine respiratory coronavirus
Porcine epidemic diarrhea virus
Feline coronavirus
Canine coronavirus
Rat coronavirus

**FIGURE S9:** A heat map depicting the hierarchal clustering of the *dS* values of the Spike protein encoding ORFs (Values within SARS-CoV2, SARS-CoV, MERS-CoV and Bat-CoV are marked with a black dot).

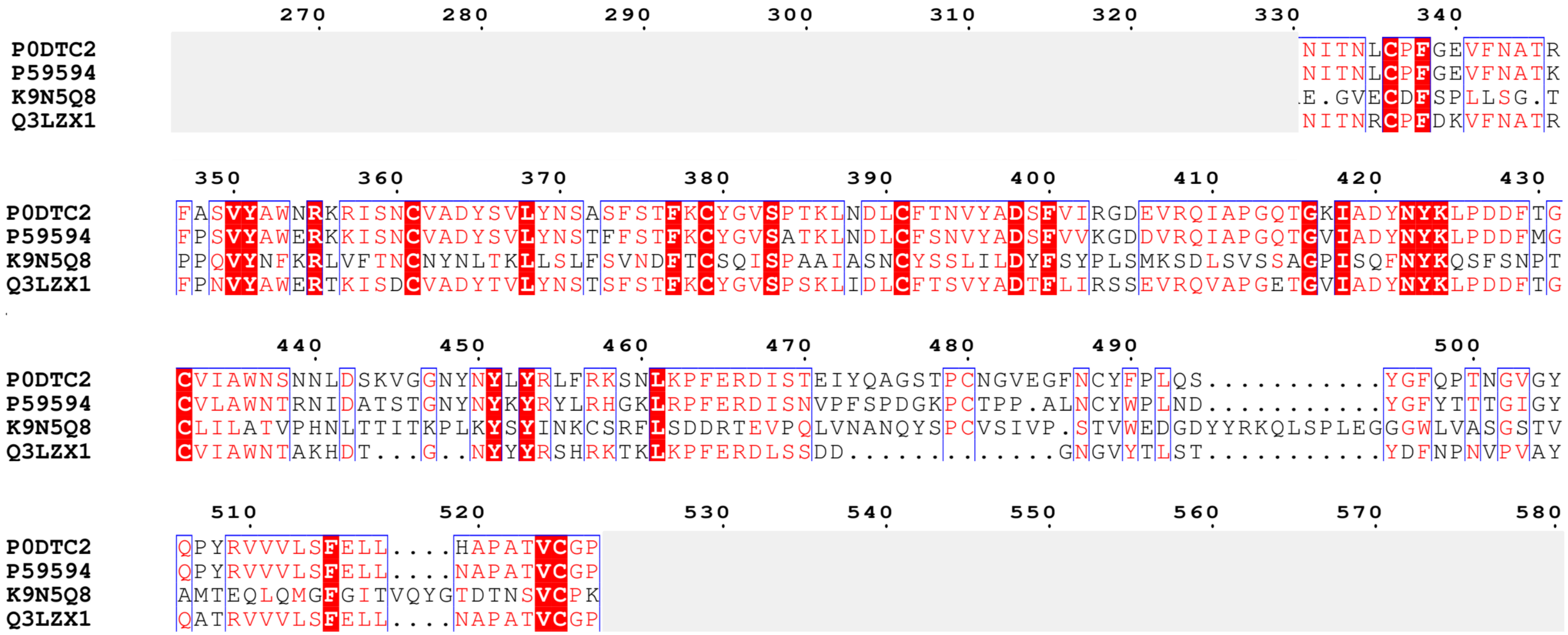
270 280 290 300 310 320 330 340
P0DTC2 NITNLCPFGEVFNATR
P59594 NITNLCPFGEVFNATK
K9N5Q8 E.GVECDFSPLLSG.T
Q3LZX1 NITNRCPFDKVFNATR
350 360 370 380 390 400 410 420 430
P0DTC2 FASVYAWNRKRISNCVADYSVLYNSASFSTFKCYGVSPTKLNDLCFTNVYADSFVIRGDEVRQIAPGQTGKIADYNYKLPDDFTG
P59594 FPSVYAWERKKISNCVADYSVLYNSTFFSTFKCYGVSATKLNDLCFSNVYADSFVVKGDDVRQIAPGQTGVIADYNYKLPDDFMG
K9N5Q8 PPQVYNFKRLVFTNCNYNLTKLLSLFSVNDFTCSQISPAAIASNCYSSLILDYFSYPLSMKSDLSVSSAGPISQFNYKQSFSNPT
Q3LZX1 FPNVYAWERTKISDCVADYTVLYNSTSFSTFKCYGVSPSKLIDLCFTSVYADTFLIRSSEVRQVAPGETGVIADYNYKLPDDFTG
440 450 460 470 480 490 500
P0DTC2 CVIAWNSNNLDSKVGGNYNYLYRLFRKSNLKPFERDISTEIYQAGSTPCNGVEGFNCYFPLQS...........YGFQPTNGVGY
P59594 CVLAWNTRNIDATSTGNYNYKYRYLRHGKLRPFERDISNVPFSPDGKPCTPP.ALNCYWPLND...........YGFYTTTGIGY
K9N5Q8 CLILATVPHNLTTITKPLKYSYINKCSRFLSDDRTEVPQLVNANQYSPCVSIVP.STVWEDGDYYRKQLSPLEGGGWLVASGSTV
Q3LZX1 CVIAWNTAKHDT...G..NYYYRSHRKTKLKPFERDLSSDD.............GNGVYTLST...........YDFNPNVPVAY
510 520 530 540 550 560 570 580
P0DTC2 QPYRVVVLSFELL....HAPATVCGP
P59594 QPYRVVVLSFELL....NAPATVCGP
K9N5Q8 AMTEQLQMGFGITVQYGTDTNSVCPK
Q3LZX1 QATRVVVLSFELL....NAPATVCGP

**FIGURE S10:** A multiple protein sequence alignment of the receptor binding CTDs of the Spike proteins from SARS-CoV, Bat-CoV, MERS-CoV and SARS-CoV2.

```
         960       970       980       990       1000      1010      1020      1030
P0DTC2                                      AATATTACAAACTTGTGCCCTTTTGGTGAAGTTTTTAACGCCACCAGA
P59594                                      AATATTACAAACTTGTGTCCTTTTGGAGAGGTTTTTAATGCTACTAAA
K9N5Q8                                      GAA...GGTGTTGAATGTGATTTTTCACCTCTTCTGTCTGGC...ACA
Q3LZX1                                      AACATTACAAATCGCTGTCCTTTTGACAAGGTTTTTAATGCTACTCGC

       1040      1050      1060      1070      1080      1090      1100      1110      1120
P0DTC2 TTTGCATCTGTTTATGCTTGGAACAGGAAGAGAATCAGCAACTGTGTTGCTGATTATTCTGTCCTATATAATTCCGCATCATTTT
P59594 TTCCCTTCTGTCTATGCATGGGAGAGAAAAAAAATTTCTAATTGTGTTGCTGATTACTCTGTGCTCTACAACTCAACATTTTTTT
K9N5Q8 CCTCCTCAGGTTTATAATTTCAAGCGTTTGGTTTTTACCAATTGCAATTATAATCTTACCAAATTGCTTTCACTTTTTTCTGTGA
Q3LZX1 TTTCCTAATGTTTATGCATGGGAGAGAACAAAAATCTCTGATTGTGTTGCTGACTACACTGTTCTCTACAACTCAACCTCTTTCT

           1130      1140      1150      1160      1170      1180      1190      1200
P0DTC2 CCACTTTTAAGTGTTATGGAGTGTCTCCTACTAAATTAAATGATCTCTGCTTTACTAATGTCTATGCAGATTCATTTGTAATTAG
P59594 CAACCTTTAAGTGCTATGGCGTTTCTGCCACTAAGTTGAATGATCTTTGCTTCTCCAATGTCTATGCAGATTCTTTTGTAGTCAA
K9N5Q8 ATGATTTTACTTGTAGTCAAATATCTCCAGCAGCAATTGCTAGTAACTGTTATTCTTCACTGATTTTGGATTATTTTCATACCC
Q3LZX1 CGACTTTTAAATGTTATGGAGTGTCTCCATCTAAGTTGATTGACCTATGCTTTACAAGTGTGTATGCTGATACATTCTTGATAAG

       1210      1220      1230      1240      1250      1260      1270      1280      1290
P0DTC2 AGGTGATGAAGTCAGACAAATCGCTCCAGGGCAAACTGGAAAGATTGCTGATTATAATTATAAATTACCAGATGATTTTACAGGC
P59594 GGGAGATGATGTAAGACAAATAGCGCCAGGACAAACTGGTGTTATTGCTGATTATAATTATAAATTGCCAGATGATTTCATGGGT
K9N5Q8 ACTTAGTATGAAATCCGATCTCAGTGTTAGTTCTGCTGGTCCAATATCCCAGTTTAATTATAAACAGTCCTTTTCTAATCCCACA
Q3LZX1 ATCTTCTGAAGTAAGACAAGTTGCACCAGGTGAAACTGGTGTTATTGCTGACTACAATTACAAGTTGCCTGATGATTTCACTGGC

           1300      1310      1320      1330      1340      1350      1360      1370
P0DTC2 TGCGTTATAGCTTGGAATTCTAACAATCTTGATTCTAAGGTTGGTGGTAATTATAATTACCTGTATAGATTGTTTAGGAAGTCTA
P59594 TGTGTCCTTGCTTGGAATACTAGGAACATTGATGCTACTTCAACTGGTAATTATAATTATAAATATAGGTATCTTAGACATGGCA
K9N5Q8 TGTTTGATTTTAGCGACTGTTCCTCATAACCTTACTACTATTACTAAGCCTCTTAAGTACAGCTATATTAACAAGTGCTCTCGTT
Q3LZX1 TGTGTAATTGCTTGGAATACTGCTAAACATGATACT.........GGC......AATTATTACTACAGATCTCATCGCAAGACTA

       1380      1390      1400      1410      1420      1430      1440      1450      1460
P0DTC2 ATCTCAAACCTTTTGAGAGAGATATTTCAACTGAAATCTATCAGGCCGGTAGCACACCTTGTAATGGTGTTGAAGGTTTTAATTG
P59594 AGCTTAGGCCCCTTTGAGAGAGACATATCTAATGTGCCTTTCTCCCCTGATGGCAAACCTTGCACCCCACCT...GCTCTTAATTG
K9N5Q8 TTCTTTCTGATGATCGTACTGAAGTACCTCAGTTAGTGAACGCTAATCAATACTCACCCTGTGTATCCATTGTCCCA...TCCAC
Q3LZX1 AGTTAAAGCCTTTTGAGAGAGACCTGTCTTCTGACGAT.......................................GGTAATGG

           1470      1480                                          1490      1500      1510
P0DTC2 TTACTTTCCTTTACAATCA.................................TATGGTTTCCAACCCACTAATGGTGTTGGTTAC
P59594 TTATTGGCCATTAAATGAT.................................TATGGTTTTTACACCACTACTGGCATTGGCTAC
K9N5Q8 TGTGTGGGAAGACGGTGATTATTATAGGAAACAACTATCTCCACTTGAAGGTGGTGGCTGGCTTGTTGCTAGTGGCTCAACTGTT
Q3LZX1 TGTGTATACACTCTCAACA.................................TATGACTTTAACCCTAACGTTCCAGTAGCATAT

          1520      1530      1540      1550                1560      1570      1580
P0DTC2 CAACCATACAGAGTAGTAGTACTTTCTTTTGAACTTCTA............CATGCACCAGCAACTGTTTGTGGACCT
P59594 CAACCTTACAGAGTTGTAGTACTTTCTTTTGAACTTTTA............AATGCACCGGCCACGGTTTGTGGACCA
K9N5Q8 GCCATGACTGAGCAATTACAGATGGGCTTTGGTATTACAGTTCAATATGGTACAGACACCAATAGTGTTTGCCCCAAG
Q3LZX1 CAGGCTACTAGGGTTGTTGTACTCTCTTTTGAACTTCTT............AATGCACCTGCTACAGTTTGTGGACCT
```

**FIGURE S11:** A multiple coding nucleotide sequence alignment of the receptor binding CTDs of the Spike proteins from SARS-CoV, Bat-CoV, MERS-CoV and SARS-CoV2.

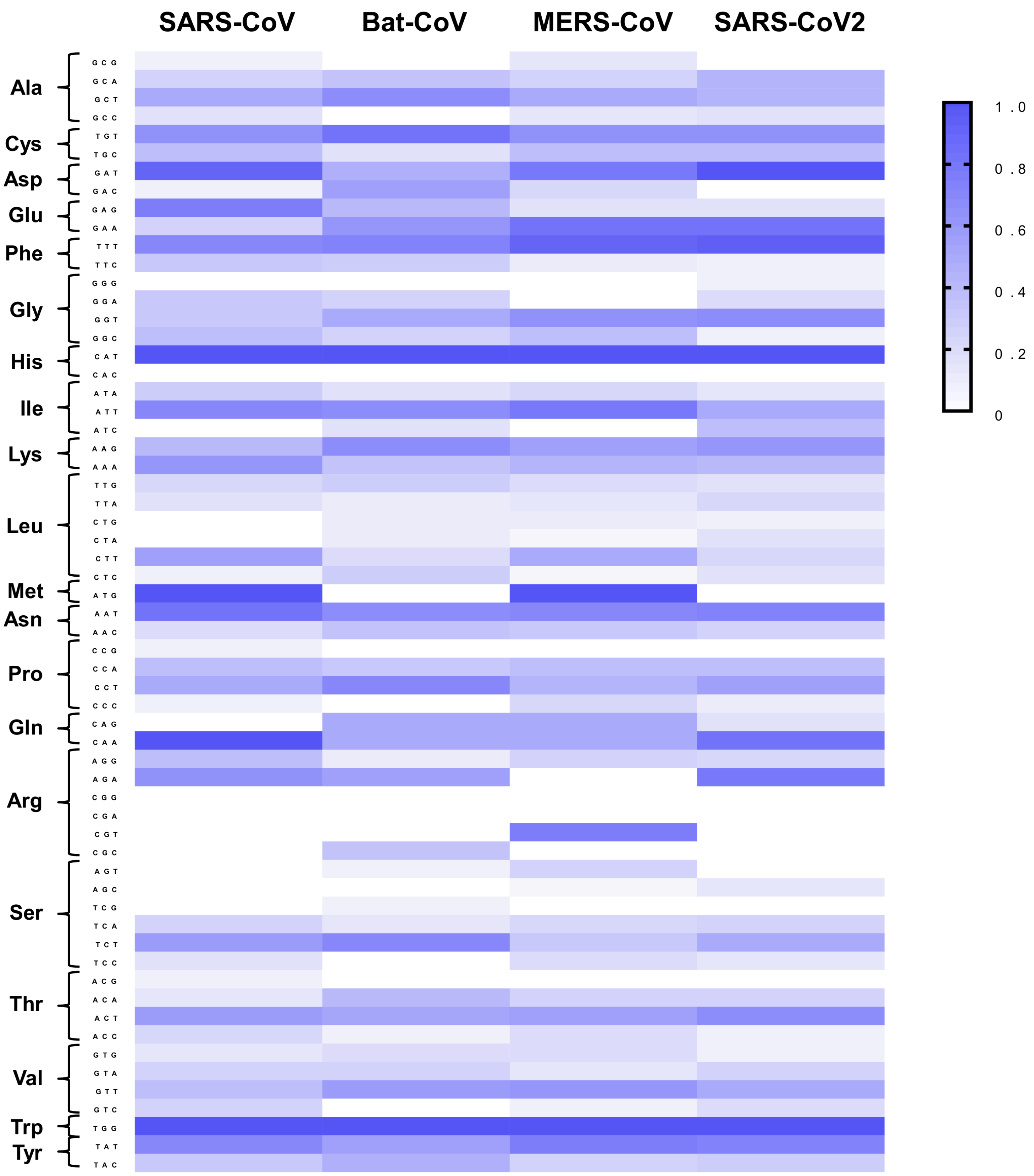
SARS-CoV
Bat-CoV
MERS-CoV
SARS-CoV2
Ala
GCG
GCA
GCT
GCC
Cys
TGT
TGC
Asp
GAT
GAC
Glu
GAG
GAA
Phe
TTT
TTC
Gly
GGG
GGA
GGT
GGC
His
CAT
CAC
Ile
ATA
ATT
ATC
Lys
AAG
AAA
Leu
TTG
TTA
CTG
CTA
CTT
CTC
Met
ATG
Asn
AAT
AAC
Pro
CCG
CCA
CCT
CCC
Gln
CAG
CAA
Arg
AGG
AGA
CGG
CGA
CGT
CGC
Ser
AGT
AGC
TCG
TCA
TCT
TCC
Thr
ACG
ACA
ACT
ACC
Val
GTG
GTA
GTT
GTC
Trp
TGG
Tyr
TAT
TAC
1.0
0.8
0.6
0.4
0.2
0

**FIGURE S12:** A comparison of the codon usage in the CTDs of the Spike proteins from SARS-CoV, Bat-CoV, MERS-CoV and SARS-CoV2.

### Supplementary Tables

#### Table S1: List of Spike Proteins' Gene (NCBI) and Protein (UniProt) accession IDs.

| Organism | Year | Gene ID | Protein ID |
|---|---|---|---|
| Severe acute respiratory syndrome coronavirus 2 (2019-nCoV) (SARS-CoV-2) | 2020 | QHD43416 | P0DTC2 |
| Middle East respiratory syndrome-related coronavirus (isolate United Kingdom/H123990006/2012) (Betacoronavirus England 1) (Human coronavirus EMC) | 2013 | AFY13307 | K9N5Q8 |
| Bat coronavirus 133/2005 (BtCoV) (BtCoV/133/2005) | 2007 | ABG47052 | Q0Q4F2 |
| Bat coronavirus HKU3 (BtCoV) (SARS-like coronavirus HKU3) | 2007 | AAY88866 | Q3LZX1 |
| Bat coronavirus HKU4 (BtCoV) (BtCoV/HKU4/2004) | 2007 | ABN10839 | A3EX94 |
| Bat coronavirus HKU5 (BtCoV) (BtCoV/HKU5/2004) | 2007 | ABN10875 | A3EXD0 |
| Bat coronavirus HKU9 (BtCoV) (BtCoV/HKU9) | 2007 | ABN10911 | A3EXG6 |
| Bovine coronavirus (strain 98TXSF-110-ENT) (BCoV-ENT) (BCV) | 2007 | AAK83356 | Q91A26 |
| Bovine coronavirus (strain 98TXSF-110-LUN) (BCoV-LUN) (BCV) | 2007 | AAL57308 | Q8V436 |
| Bovine coronavirus (strain OK-0514) (BCoV) (BCV) | 2007 | AAF25519 | Q9QAQ8 |
| Canine coronavirus (strain BGF10) (CCoV) (Canine enteric coronavirus) | 2007 | AAQ17220 | Q7T6T3 |
| Human coronavirus HKU1 (isolate N1) (HCoV-HKU1) | 2007 | AAT98580 | Q5MQD0 |
| Porcine epidemic diarrhea virus (strain CV777) (PEDV) | 2007 | AAK38656 | Q91AV1 |
| Porcine hemagglutinating encephalomyelitis virus (strain 67N) (HEV-67N) | 2007 | AAL80031 | Q8BB25 |
| Porcine hemagglutinating encephalomyelitis virus (strain IAF-404) (HEV) | 2007 | AAM77000 | Q8JSP8 |
| Rat coronavirus (strain 681) (RCV-SDAV) (Sialodacryoadenitis virus SDAV-681) | 2007 | AAF97738 | Q9IKD1 |
| Bat coronavirus Rp3/2004 (BtCoV/Rp3/2004) (SARS-like coronavirus Rp3) | 2005 | AAZ67052 | Q3I5J5 |
| Canine coronavirus (strain K378) (CCoV) (Canine enteric coronavirus) | 2005 | CAA54335 | Q65984 |
| Human coronavirus NL63 (HCoV-NL63) | 2005 | AAS89767 | Q6Q1S2 |

| | | | |
|---|---|---|---|
| Human SARS coronavirus (SARS-CoV) (Severe acute respiratory syndrome coronavirus) | 2003 | AAP13441 | P59594 |
| Canine coronavirus (strain Insavc-1) (CCoV) (Canine enteric coronavirus) | 1994 | BAA02408 | P36300 |
| Human coronavirus OC43 (HCoV-OC43) | 1994 | AAA03055 | P36334 |
| Murine coronavirus (strain JHMV / variant CL-2) (MHV) (Murine hepatitis virus) | 1994 | BAA01085 | Q02385 |
| Porcine transmissible gastroenteritis coronavirus (strain Miller) (TGEV) | 1994 | AAB19567 | P33470 |
| Porcine transmissible gastroenteritis coronavirus (strain NEB72-rt) (TGEV) | 1994 | AAA47108 | Q01977 |
| Bovine coronavirus (strain F15) (BCoV) (BCV) | 1992 | BAA00631 | P25190 |
| Bovine coronavirus (strain L9) (BCoV) (BCV) | 1992 | AAA42907 | P25191 |
| Bovine coronavirus (strain LY-138) (BCoV) (BCV) | 1992 | AAF25499 | P25192 |
| Bovine coronavirus (strain Quebec) (BCoV) (BCV) | 1992 | AAL40400 | P25193 |
| Bovine coronavirus (strain vaccine) (BCoV) (BCV) | 1992 | AAA42908 | P25194 |
| Porcine respiratory coronavirus (strain 86/137004 / isolate British) (PRCoV) (PRCV) | 1992 | CAA42686 | P27655 |
| Porcine respiratory coronavirus (strain RM4) (PRCoV) (PRCV) | 1992 | CAA80837 | P24413 |
| Murine coronavirus (strain 4) (MHV-4) (Murine hepatitis virus) | 1991 | AAA46456 | P22432 |
| Bovine coronavirus (strain Mebus) (BCoV) (BCV) | 1990 | AAA66399 | P15777 |
| Human coronavirus 229E (HCoV-229E) | 1990 | AAG48592 | P15423 |
| Porcine transmissible gastroenteritis coronavirus (strain FS772/70) (TGEV) | 1990 | CAA37285 | P18450 |
| Avian infectious bronchitis virus (strain Beaudette) (IBV) | 1989 | AAY21248 | P11223 |
| Avian infectious bronchitis virus (strain D274) (IBV) | 1989 | CAA33837 | P12722 |
| Avian infectious bronchitis virus (strain KB8523) (IBV) | 1989 | AAA66578 | P12650 |
| Avian infectious bronchitis virus (strain M41) (IBV) | 1989 | ABI26423 | P12651 |
| Feline coronavirus (strain FIPV WSU-79/1146) (FCoV) | 1989 | AAY16375 | P10033 |
| Murine coronavirus (strain A59) (MHV-A59) (Murine hepatitis virus) | 1989 | AAB86819 | P11224 |
| Murine coronavirus (strain JHM) (MHV-JHM) (Murine hepatitis virus) | 1989 | CAA28484 | P11225 |
| Porcine transmissible gastroenteritis coronavirus (strain Purdue) (TGEV) | 1988 | CAB91145 | P07946 |

| | | | |
|---|---|---|---|
| Avian infectious bronchitis virus (strain 6/82) (IBV) | 1987 | CAA28432 | P05135 |

**Table S2: Comparison of the amino acids and respective codons form the Subunit S1 C-Terminal domains/Receptor binding domains of Coronavirus Spike proteins.**

| Position (wrt CoV2) | Codon | | | | Amino Acids | | | |
|---|---|---|---|---|---|---|---|---|
| | CoV | Bat | MERS | CoV2 | CoV | Bat | MERS | CoV2 |
| 331 | AAT | AAC | GAA | AAT | N | N | E | N |
| 332 | ATT | ATT | --- | ATT | I | I | --- | I |
| 333 | ACA | ACA | GGT | ACA | T | T | G | T |
| 334 | AAC | AAT | GTT | AAC | N | N | V | N |
| 335 | TTG | CGC | GAA | TTG | L | E | R | L |
| 337 | CCT | CCT | GAT | CCT | P | P | D | P |
| 339 | GGA | GAC | TCA | GGT | G | D | S | G |
| 340 | GAG | AAG | CCT | GAA | E | K | P | E |
| 341 | GTT | GTT | CTT | GTT | V | V | L | V |
| 342 | TTT | TTT | CTG | TTT | F | F | L | F |
| 343 | AAT | AAT | TCT | AAC | N | N | S | N |
| 344 | GCT | GCT | GGC | GCC | A | A | G | A |
| 345 | ACT | ACT | --- | ACC | T | T | --- | T |
| 346 | AAA | CGC | ACA | AGA | R | R | T | K |
| 347 | TTC | TTT | CCT | TTT | F | F | P | F |
| 348 | CCT | CCT | CCT | GCA | P | P | P | A |
| 349 | TCT | AAT | CAG | TCT | S | N | Q | S |
| 352 | GCA | GCA | AAT | GCT | A | A | N | A |
| 353 | TGG | TGG | TTC | TGG | W | W | F | W |
| 354 | GAG | GAG | AAG | AAC | E | E | K | N |
| 356 | AAA | ACA | TTG | AAG | K | T | L | K |
| 357 | AAA | AAA | GTT | AGA | K | K | V | R |
| 358 | ATT | ATC | TTT | ATC | I | I | F | I |
| 359 | TCT | TCT | ACC | AGC | S | S | T | S |
| 360 | AAT | GAT | AAT | AAC | N | D | N | N |

| | | | | | | | | |
|---|---|---|---|---|---|---|---|---|
| 362 | GTT | GTT | AAT | GTT | V | V | N | V |
| 363 | GCT | GCT | TAT | GCT | A | A | Y | A |
| 364 | GAT | GAC | AAT | GAT | D | D | N | D |
| 365 | TAC | TAC | CTT | TAT | Y | Y | L | Y |
| 366 | TCT | ACT | ACC | TCT | S | T | T | S |
| 367 | GTG | GTT | AAA | GTC | V | V | K | V |
| 369 | TAC | TAC | CTT | TAT | Y | Y | L | Y |
| 370 | AAC | AAC | TCA | AAT | N | N | S | N |
| 371 | TCA | TCA | CTT | TCC | S | S | L | S |
| 372 | ACA | ACC | TTT | GCA | T | T | F | A |
| 373 | TTT | TCT | TCT | TCA | F | S | S | S |
| 374 | TTT | TTC | GTG | TTT | F | F | V | F |
| 375 | TCA | TCG | AAT | TCC | S | S | N | S |
| 376 | ACC | ACT | GAT | ACT | T | T | D | T |
| 378 | AAG | AAA | ACT | AAG | K | K | T | K |
| 380 | TAT | TAT | AGT | TAT | Y | Y | S | Y |
| 381 | GGC | GGA | CAA | GGA | G | G | Q | G |
| 382 | GTT | GTG | ATA | GTG | V | V | I | V |
| 384 | GCC | CCA | CCA | CCT | A | P | P | P |
| 385 | ACT | TCT | GCA | ACT | T | S | A | T |
| 386 | AAG | AAG | GCA | AAA | K | K | A | K |
| 387 | TTG | TTG | ATT | TTA | L | L | I | L |
| 388 | AAT | ATT | GCT | AAT | N | I | A | N |
| 389 | GAT | GAC | AGT | GAT | D | D | S | D |
| 390 | CTT | CTA | AAC | CTC | L | L | N | L |
| 392 | TTC | TTT | TAT | TTT | F | F | Y | F |
| 393 | TCC | ACA | TCT | ACT | S | T | S | T |
| 394 | AAT | AGT | TCA | AAT | N | S | S | N |
| 395 | GTC | GTG | CTG | GTC | V | V | L | V |
| 396 | TAT | TAT | ATT | TAT | Y | Y | I | Y |
| 397 | GCA | GCT | TTG | GCA | A | D | L | A |
| 399 | TCT | ACA | TAT | TCA | S | T | Y | S |

| | | | | | | | | |
|---|---|---|---|---|---|---|---|---|
| 401 | GTA | TTG | TCA | GTA | V | V | S | V |
| 402 | GTC | ATA | TAC | ATT | V | I | Y | I |
| 403 | AAG | AGA | CCA | AGA | K | R | P | R |
| 404 | GGA | TCT | CTT | GGT | G | S | L | G |
| 405 | GAT | TCT | AGT | GAT | D | S | S | D |
| 406 | GAT | GAA | ATG | GAA | D | E | M | E |
| 407 | GTA | GTA | AAA | GTC | V | V | K | V |
| 408 | AGA | AGA | TCC | AGA | R | R | S | R |
| 409 | CAA | CAA | GAT | CAA | Q | Q | D | Q |
| 410 | ATA | GTT | CTC | ATC | I | V | L | I |
| 411 | GCG | GCA | AGT | GCT | A | A | S | A |
| 412 | CCA | CCA | GTT | CCA | P | P | V | P |
| 413 | GGA | GGT | AGT | GGG | G | G | S | G |
| 414 | CAA | GAA | TCT | CAA | Q | E | S | Q |
| 415 | ACT | ACT | GCT | ACT | T | T | A | T |
| 417 | GTT | GTT | CCA | AAG | V | V | P | K |
| 419 | GCT | GCT | TCC | GCT | A | A | S | A |
| 420 | GAT | GAC | CAG | GAT | D | D | Q | D |
| 421 | TAT | TAC | TTT | TAT | Y | Y | F | Y |
| 425 | TTG | TTG | CAG | TTA | L | L | Q | L |
| 426 | CCA | CCT | TCC | CCA | P | P | S | P |
| 427 | GAT | GAT | TTT | GAT | D | D | F | D |
| 428 | GAT | GAT | TCT | GAT | D | D | S | D |
| 429 | TTC | TTC | AAT | TTT | F | F | N | F |
| 430 | ATG | ACT | CCC | ACA | M | T | P | T |
| 431 | GGT | GGC | ACA | GGC | G | G | T | G |
| 433 | GTC | GTA | TTG | GTT | V | V | L | V |
| 434 | CTT | ATT | ATT | ATA | L | I | I | I |
| 435 | GCT | GCT | TTA | GCT | A | A | L | A |
| 436 | TGG | TGG | GCG | TGG | W | W | A | W |
| 437 | AAT | AAT | ACT | AAT | N | N | T | N |
| 438 | ACT | ACT | GTT | TCT | T | T | V | S |

| | | | | | | | | |
|---|---|---|---|---|---|---|---|---|
| 439 | AGG | GCT | CCT | AAC | R | A | P | N |
| 440 | AAC | AAA | CAT | AAT | N | K | H | N |
| 441 | ATT | CAT | AAC | CTT | I | H | N | L |
| 442 | GAT | GAT | CTT | GAT | D | D | L | D |
| 443 | GCT | ACT | ACT | TCT | A | T | T | S |
| 444 | ACT | --- | ACT | AAG | T | --- | T | K |
| 445 | TCA | --- | ATT | GTT | S | --- | I | V |
| 446 | ACT | --- | ACT | GGT | T | --- | T | G |
| 447 | GGT | GGC | AAG | GGT | G | G | K | G |
| 448 | AAT | --- | CCT | AAT | N | --- | P | N |
| 449 | TAT | --- | CTT | TAT | Y | --- | L | Y |
| 450 | AAT | AAT | AAG | AAT | N | N | K | N |
| 452 | AAA | TAC | AGC | CTG | K | Y | S | L |
| 454 | AGG | AGA | ATT | AGA | R | R | I | R |
| 455 | TAT | TCT | AAC | TTG | Y | S | N | L |
| 456 | CTT | CAT | AAG | TTT | L | H | K | F |
| 457 | AGA | CGC | TGC | AGG | R | R | C | R |
| 458 | CAT | AAG | TCT | AAG | H | K | S | K |
| 459 | GGC | ACT | CGT | TCT | G | T | R | S |
| 460 | AAG | AAG | TTT | AAT | K | K | F | N |
| 462 | AGG | AAG | TCT | AAA | R | K | S | K |
| 463 | CCC | CCT | GAT | CCT | P | P | D | P |
| 464 | TTT | TTT | GAT | TTT | F | F | D | F |
| 465 | GAG | GAG | CGT | GAG | E | E | R | E |
| 466 | AGA | AGA | ACT | AGA | R | R | T | R |
| 467 | GAC | GAC | GAA | GAT | D | D | E | D |
| 468 | ATA | CTG | GTA | ATT | I | L | V | I |
| 469 | TCT | TCT | CCT | TCA | S | S | P | S |
| 470 | AAT | TCT | CAG | ACT | N | S | Q | T |
| 471 | GTG | GAC | TTA | GAA | V | D | L | E |
| 472 | CCT | GAT | GTG | ATC | P | D | V | I |
| 473 | TTC | --- | AAC | TAT | F | --- | N | Y |

| | | | | | | | | |
|---|---|---|---|---|---|---|---|---|
| 474 | TCC | --- | GCT | CAG | S | --- | A | Q |
| 475 | CCT | --- | AAT | GCC | P | --- | N | A |
| 476 | GAT | --- | CAA | GGT | D | --- | Q | G |
| 477 | GGC | --- | TAC | AGC | G | --- | Y | S |
| 478 | AAA | --- | TCA | ACA | K | --- | S | T |
| 479 | CCT | --- | CCC | CCT | P | --- | P | P |
| 480 | TGC | --- | TGT | TGT | C | --- | C | C |
| 481 | ACC | --- | GTA | AAT | T | --- | V | N |
| 482 | CCA | --- | TCC | GGT | P | --- | S | G |
| 483 | CCT | --- | ATT | GTT | P | --- | I | V |
| 484 | --- | --- | GTC | GAA | --- | --- | V | E |
| 485 | GCT | --- | CCA | GGT | A | --- | P | G |
| 486 | CTT | GGT | --- | TTT | L | G | --- | F |
| 487 | AAT | AAT | TCC | AAT | N | N | S | N |
| 488 | TGT | GGT | ACT | TGT | C | G | T | C |
| 489 | TAT | GTG | GTG | TAC | Y | V | V | Y |
| 490 | TGG | TAT | TGG | TTT | W | Y | W | F |
| 491 | CCA | ACA | GAA | CCT | P | T | E | P |
| 492 | TTA | CTC | GAC | TTA | L | L | D | L |
| 493 | AAT | TCA | GGT | CAA | N | S | G | Q |
| 494 | GAT | ACA | GAT | TCA | D | T | D | S |
| 495 | TAT | TAT | GGT | TAT | Y | Y | G | Y |
| 496 | GGT | GAC | GGC | GGT | G | D | G | G |
| 497 | TTT | TTT | TGG | TTC | F | F | W | F |
| 498 | TAC | AAC | CTT | CAA | Y | N | N | Q |
| 499 | ACC | CCT | GTT | CCC | T | P | V | P |
| 500 | ACT | AAC | GCT | ACT | T | N | A | T |
| 501 | ACT | GTT | AGT | AAT | T | V | S | N |
| 502 | GGC | CCA | GGC | GGT | G | P | G | G |
| 503 | ATT | GTA | TCA | GTT | I | V | S | V |
| 504 | GGC | GCA | ACT | GGT | G | A | T | G |
| 505 | TAC | TAT | GTT | TAC | Y | Y | V | Y |

| | | | | | | | | |
|---|---|---|---|---|---|---|---|---|
| 506 | CAA | CAG | GCC | CAA | Q | Q | A | Q |
| 507 | CCT | GCT | ATG | CCA | P | A | M | P |
| 508 | TAC | ACT | ACT | TAC | Y | T | T | Y |
| 509 | AGA | AGG | GAG | AGA | R | R | E | R |
| 510 | GTT | GTT | CAA | GTA | V | V | Q | V |
| 511 | GTA | GTT | TTA | GTA | V | V | L | V |
| 512 | GTA | GTA | CAG | GTA | V | V | Q | V |
| 513 | CTT | CTC | ATG | CTT | L | L | M | L |
| 514 | TCT | TCT | GGC | TCT | S | S | G | S |
| 516 | GAA | GAA | GGT | GAA | E | E | G | E |
| 517 | CTT | CTT | ATT | CTT | L | L | I | L |
| 518 | TTA | CTT | ACA | CTA | L | L | T | L |
| 519 | AAT | AAT | ACA | CAT | N | N | T | H |
| 520 | GCA | GCA | GAC | GCA | A | A | D | A |
| 521 | CCG | CCT | ACC | CCA | P | P | T | P |
| 522 | GCC | GCT | AAT | GCA | A | A | N | A |
| 523 | ACG | ACA | AGT | ACT | T | T | S | T |
| 526 | GGA | GGA | CCC | GGA | G | G | P | G |
| 527 | CCA | CCT | AAG | CCT | P | P | K | P |
| **% of charged amino acids (D, E, K, R)** | | | | | 17.3 | 20.1 | 13.6 | 16.3 |
| **% of aromatic amino acids (F, Y, W)** | | | | | 16.3 | 14.5 | 14.6 | 15.7 |
| **Grand average of hydropathicity** | | | | | -0.22 | -0.26 | -0.067 | -0.216 |
| **Calculated pI** | | | | | 8.1 | 6.89 | 5.08 | 8.11 |
| --- indicates gaps | | | | | | | | |
